# Novel Herpesvirus Transcripts with Putative Regulatory Roles in DNA Replication and Global Transcription

**DOI:** 10.1101/2023.03.25.534217

**Authors:** Gábor Torma, Dóra Tombácz, Islam A.A. Almsarrhad, Zsolt Csabai, Gergely Ármin Nagy, Balázs Kakuk, Gábor Gulyás, Lauren McKenzie Spires, Ishaan Gupta, Ádám Fülöp, Ákos Dörmő, István Prazsák, Máté Mizik, Virág Éva Dani, Viktor Csányi, Zoltán Zádori, Zsolt Toth, Zsolt Boldogkői

## Abstract

In the last couple of years, the rapid advances and decreasing costs of sequencing technologies have revolutionized transcriptomic research. Long-read sequencing (LRS) techniques are able to detect full-length RNA molecules in a single run without the need for additional assembly steps. LRS studies have revealed an unexpected transcriptomic complexity in a variety of organisms, including viruses. A number of transcripts with proven or putative regulatory role, mapping close to or overlapping the replication origins (Oris) and the nearby transcription activator genes, have been described in herpesviruses. In this study, we applied both newly generated and previously published LRS and short-read sequencing datasets to discover additional Ori-proximal transcripts in nine herpesviruses belonging to all of the three subfamilies (alpha, beta and gamma). We identified novel long non-coding RNAs (lncRNAs), as well as splice and length isoforms of mRNAs and lncRNAs. Furthermore, our analysis disclosed an intricate meshwork of transcriptional overlaps at the examined genomic regions. Our results suggest the existence of a ‘super regulatory center’, which controls both the replication and the global transcription through multilevel interactions between the molecular machineries.

## INTRODUCTION

### Regulation of herpesvirus transcription

The expression of herpesvirus genes is complexly regulated by both cellular and viral factors during productive infection. Four immediate early (IE) proteins are involved in the transcription regulation of herpes simplex virus type 1 (HSV-1), a representative member of alphaherpesviruses (αHVs). The essential ICP4 viral protein [encoded by *rs1* (*icp4*)] is the main transcription regulator of HSV-1, which recruits cellular transcription factors (TFs; e.g. TFIID) to viral promoters in order to enhance (or in some cases to repress) transcription initiation^1^. ICP22 (encoded by *us1*) has been shown to promote the transcription elongation of viral RNAs^2^. The *us1* gene of HSV-1 is located in the unique short (US) genomic region in a single copy while its promoter is located in duplicate in the inverted repeat (IR) region (the other IR copy controls the expression of *ul12* gene). In the Varicellovirus genus, the *us1* gene is translocated to the IR region where it is present in duplicate. ICP0 (encoded by *rl2* [*icp0*)], in a strict sense, is not a TF because it does not bind DNA or other TFs. This viral protein can promote viral gene expression through affecting prechromatin interactions before histones are bound the viral DNA^3^. Similar to the viral proteins mentioned above, ICP27 (encoded by *ul54*) is also multifunctional. It is involved in recruiting the RNA polymerase (RNP) to viral promoters^4^, and also in the post-transcriptional regulation of gene expression and the synthesis of viral DNA^5^.

The genetic control of the lytic cycle of other αHVs is very similar to that of HSV-1. One of the major differences is the expression kinetics of the *rl2*, *us1* and *ul54* orthologs: they have evolved to be expressed in early (E) kinetics in pseudorabies virus (PRV) and equid alphaherpesvirus type 1 (EHV-1). Additionally, the *icp0* gene has been structurally and functionally simplified in these viruses. The betaherpesviruses (βHVs), and gammaherpesviruses (γHVs) utilize basically similar mechanisms as those of αHVs for the control of genome-wide viral transcription. In human cytomegalovirus (HCMV), the prototype member of βHVs, two major IE genes (*ie1* and *ie2*) regulate the global viral transcription^6^. The IE proteins of Epstein-Barr virus (EBV; the representative member of HVs), designated as BZLF1 and BRLF1, are transactivators that turn on the transcription of E viral genes^7^.

The latency associated transcript (LAT) of HSV-1 is the only viral gene that is highly expressed during the latent state^8^. This non-coding RNA transcript represses the expression of lytic viral genes through blocking ICP4 expression^9^ and facilitating heterochromatin formation on the HSV-1 genome^10^. Although LAT contains numerous open reading frames (ORFs), it likely does not encode any polypeptides^11, 12^. Other long non-coding RNAs (lncRNAs) can also be expressed during latency such as the long-latency transcript (LLT; overlapping both the *icp0* and *icp4* genes^13^), and the L/S junction-spanning transcripts (L/STs; overlapping the *icp34.5* and *icp4* genes^14^).

The highly abundant non-coding NOIR-1 transcript family described in αHVs [PRV^15^, the varicella-zoster virus (VZV)^16^, EHV-1^17^] are 3′-coterminal with the LLT/AST transcripts and they are expressed in the lytic cycle. The low-abundance NOIR-2 RNA, which is transcribed in a reverse orientation relative to NOIR-1, has only been detected in PRV^15^. ELIE, another lytic ncRNA of PRV^18^, is partially homologous to the long isoform of HSV-1 L/ST expressed during latency. The precise function of NOIR and ELIE transcripts remains to be determined.

### DNA replication

Although the mechanism of DNA replication substantially differs between the domains of life, they also share many similarities with each other. While the prokaryotic genomes contain a single start site of DNA synthesis (designated as replication origin; Ori) that is defined by consensus sequences^19^, the eukaryotic genomes typically have tens of thousands Oris that are specified by their chromatin structure^20, 21^. Viruses have a single or a few Oris, which are specified by a combination of structural properties and sequence specificity (typically AT-rich regions) of the particular DNA segment^22^. The replication of eukaryotic genomes is initiated by the binding of origin recognition complex (ORC) to the Ori^23^. The function of ORC is to serve as a platform for the assembly of the replisome, which is composed of a wide range of proteins such as the DNA helicase, DNA polymerase (DNP), topoisomerase, primase, DNA gyrase, single-stranded DNA-binding protein (ssDBP), RNase H, DNA ligase, and telomerase enzymes.

Herpesviruses encode several replication proteins needed for the DNA synthesis. For example, HSV-1 codes for an origin-binding protein (OBP) (*ul9*), an ssDBP (*ul29*), two DNPs (*ul30* and *ul42*), and three helicase/primase molecules (*ul5*, *ul8*, and *ul52*)^24, 25^. Several viral auxiliary factors play roles in the nucleotide metabolism [ribonucleotide reductase (*ul39/40*), thymidine kinase (*ul23*), uracil glycosylase (*ul12*), deoxyuridine triphosphatase (*ul50*), alkaline nuclease (*ul12*)], which allows the replication of herpesviruses in non-dividing cells^26^.

HSV-1 contains three Oris, two within the inverted repeats (IRs) surrounding the US region (OriS), and one in the unique long (UL) region (OriL) (**Supplementary Figure 1**). The OriL is located between two E genes (*ul29* and *ul30*) that play essential roles in DNA replication while the 2 copies of OriS are surrounded by the IE genes *icp4* and *us1* in the internal repeat of the US region (IRS) and by *icp4* and *us12* genes within the terminal repeat of the US region (TRS). The Oris in other Simplexviruses such as HSV-2^27^ and bovine alphaherpesvirus 2 (BoHV-2^28^) are mapped in the same genomic loci.

In contrast to the Simplexvirus genus, some members of Varicelloviruses [such as VZV, BoHV-1, and felid alphaherpesvirus 1 (FHV-1)] lack OriL. In other Varicelloviruses [such as in PRV, EHV-1, EHV-4^29^, and canid alphaherpesvirus 1 [CHV-1^30^], the position of OriL has been relocated to the intergenic region between the *ul21*-*ul22* gene pair. HCMV has similar genomic organization to HSV-1, however, it has only a single replication origin (OriLyt), which is located at similar position (next to *ul57* orthologous to the HSV-1 *ul29*). In the case of human gammaherpesviruses, EBV and KSHV have two lytic replication origins (OriLyt-L and OriLyt-R) while the latent replication is mediated by the latent replication origin (OriP) in EBV and by the terminal repeat (TR) in KSHV ^31–34^.

### Overlapping viral transcripts

Herpesviruses have a compact genome ranging in size from about 120 to 240 kb and containing 60 to 200 protein-coding genes and several ncRNAs. Recent studies^35^ have demonstrated that every viral gene forms various transcriptional overlaps (TOs), including divergent (head-to-head), convergent (tail-to-tail), and parallel (tail-to-head) TOs. The 3’-coterminal tandem genes form parallel-overlapping multigenic transcripts, which represent the archetypal genomic organization of herpesviruses. Additionally, viral genes code for 5’-truncated transcripts with ‘nested’ open reading frames (ORFs), which possess different transcription start site (TSS), but the same transcription end site (TES)^36, 37^. Most co-located divergent genes produce ‘hard’ TOs where the canonical transcripts overlap each other. In a few cases, only the long transcript isoforms (TIs), but not the canonical transcript, produce head-to head TOs (‘soft’ TOs). Convergently oriented genes form TOs through transcriptional read-through (‘soft’ TOs), but only a few ‘hard’ TOs can be observed in these gene pairs, for example in αHVs the ul7/8, *ul30/31*, and *ul50/51* gene pairs^38^.

### Non-coding RNAs regulating DNA replication

Increasing evidence has been emerging that certain non-coding transcripts, including short ncRNAs [sncRNAs, such as microRNAs (miRNAs^39^)], and lncRNAs, play crucial roles in the regulation of DNA replication^40^. The RNA primers generated by the primase enzyme and the RNA component of the telomerase^41^ have been described decades ago. Besides these, many other ncRNA classes have been discovered since then. A special type of lncRNAs encoded by the DNA sequences near the Oris have been described in all of the three domains of life and also in viruses in the past decade. A recent survey has revealed that approximately 72% of the mammalian ORC1s are associated with active promoters, 53.5% of which controls ncRNA^42^. Replication RNAs have several modes for controlling the duplication process of the DNA molecule, which includes the regulation of RNA primer synthesis through for example hybridizing with DNA sequences^43^. These RNAs can also form hybrids with mRNAs, which initiates the degradation by RNase H, thereby inhibiting the translation of replication proteins. These RNAs can also act to recruit ORC to the Ori^44^. Such transcripts have also been detected in viruses. For example, a small replication RNA has been described in human BK polyomavirus^45^. This RNA molecule binds to both DNA strands within the Ori region and inhibits the replication through interfering with the RNA primer synthesis.

### Herpesvirus transcripts mapped in the proximity of replication origins of viral genomes

Replication-origin associated RNAs (raRNAs) have been identified in all of the three subfamilies of herpesviruses. While many of these transcripts have been formerly described in βHVs and γHVs, they were practically undetected in αHVs until recently. The non-coding RNA4.9 of HCMV transcribed from the OriLyt has been identified^46^ and functionally characterized^47^. It was demonstrated that the RNA4.9 transcript regulates viral DNA replication through the formation of DNA:RNA hybrid and also has an effect on the expression level of ssDBP encoded by the *ul57* gene. RNA4.9 is supposed to have additional roles operating in both *cis* and *trans*, for example, through promoting transcriptional repression of the major IE promoter during latency^48^. The discovery of RNA4.9 and other HCMV raRNAs led to the belief that this virus has a unique mode of replication regulation, which has not proven to be true. Another non-coding replicator transcript, the SRT (smallest replicator transcript; mapping to the OriLyt) has been shown to be also essential in the DNA replication of HCMV^49^. Other replication RNAs (vRNA-1 and vRNA-2) overlapping the OriLyt have also been described in HCMV^50^.

The BHLF1 non-coding replication RNA forming an RNA:DNA hybrid at the EBV OriLyt region has been described by Rennekamp and Lieberman^51^. A bidirectional promoter within the EBV OriLyt has also been reported^52^. A highly structured RNA of EBV has also been identified^44^. The function of this transcript is to assist the viral EBNA1 and HMGA1a proteins for recruiting ORC^44^. Incomplete sequences of two co-terminal lncRNAs near the HSV-1 OriS have been described previously^53^. The emergence of long-read sequencing (LRS) technologies had a great impact on transcriptome research and accelerated the discovery of novel viral transcripts and their TIs, including splice, TSS and TES variants. These investigations have identified a number of lncRNAs near both the OriS and OriL regions of αHVs^35, 54–56^. However, since no precise function has been linked to these transcripts until now, we cannot term them as ‘replication RNAs’ that would refer to a still undisclosed role in DNA replication. Furthermore, the genes around the Oris have been shown to produce TIs with long 5′-untranslated regions (5′ UTRs – TSS isoforms), or 3′ UTRs (TES isoforms) that overlap the replication origin^57, 58^. Many or all of the mRNAs with very long 5′ UTR are probably also lncRNAs because their translation initiation sites are too far from their TSSs.

## RESULTS

### 1. Multiplatform sequencing for annotation of viral transcripts

In this work, novel and previously published sequencing datasets were used for the transcriptomic analysis of five human and four animal pathogenic herpesviruses, which were as follows: six alphaherpesviruses: [a Simplexvirus: HSV-1^57^ and five Varicelloviruses: PRV^59^; VZV^16^; BoHV-1^56^; EHV-1^17^; and simian varicella virus (SVV^60^)]; as well as a βHV [HCMV^61^]; and two γHVs [EBV^62, 63^ and Kaposi’s sarcoma-associated herpesvirus (KSHV)]. Novel sequencing was carried out by using direct cDNA sequencing (dcDNA-Seq) on MinION platform of Oxford Nanopore Technologies (ONT) for PRV, EHV-1 and KSHV, direct RNA sequencing (dRNA-Seq) for EHV-1 and KSHV, as well as amplified cDNA sequencing for VZV also on ONT device, and Illumina short-read sequencing (SRS) for EHV-1. Additionally, Cap Analysis of Gene Expression (CAGE) sequencing (CAGE-Seq, running on Illumina platform) was used for defining the TSS landscape in EHV-1 and KSHV (**Figure 1**). We also used a Terminator enzyme-based library preparation method for both dcDNA-Seq and dRNA-Seq in order to enrich the capped transcripts. Besides the oligo(dT) primer-based RT used for the techniques above, we also applied random hexamer priming for VZV sequencing. Previous data on herpesvirus transcriptomes that we reanalyzed in our study were produced by others or by our research group and they were based on the following methods (**Table 1**): SRS on various Illumina platforms^54, 64–67^, as well as LRS on ONT (MinION^17^), Pacific Biosciences (PacBio – RSII and Sequel^15, 62^), and LoopSeq^56^ platforms using a wide range of library preparation techniques and CAGE-Seq for VZV^60^ and EBV^62^.

**Figure 1.**
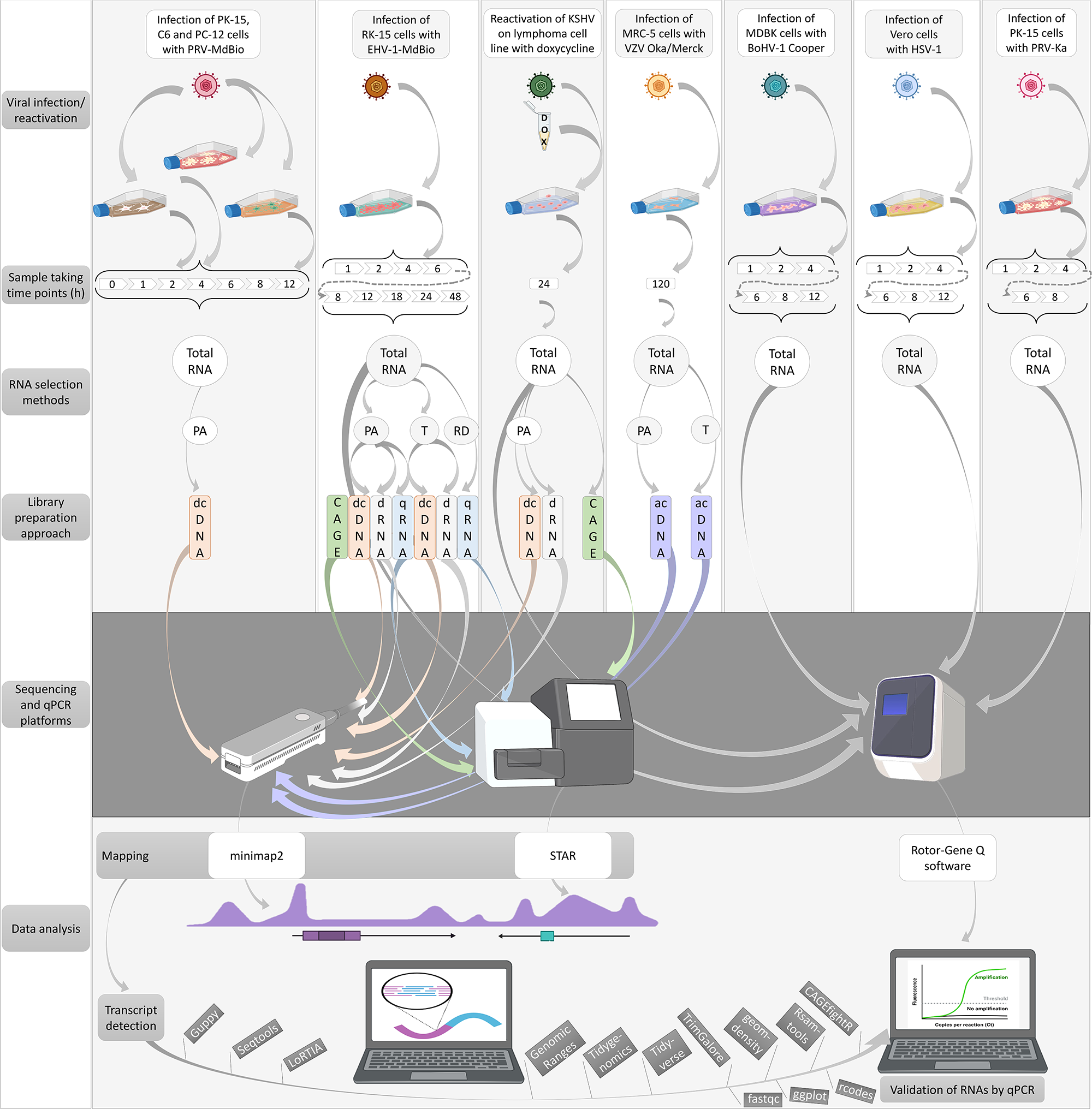
Workflow. Techniques applied for the generation of novel sequencing data, including infection of the cells with various viruses, library preparation, sequencing and bioinformatics. The workflow of the qPCR validations of several transcripts is also illustrated. Abbreviations: PA: polyadenylated RNAs; T: Terminator-handled samples; RD: ribodepleted RNAs; dcDNA: direct cDNA-seq; CAGE: Cap Analysis Gene Expression-Seq; dRNA: direct RNA-Seq; qRNA: short-read RNA-Seq (library generated by qRNA-seq kit); acDNA: amplified cDNA-Seq.

**Table 1.** Techniques and datasets of previous publications used for the analysis. This table show the annotated transcripts mapping to the genomic loci examined in this study.

| Virus | Sequencing approach | Library | Reference | Data availability |
| --- | --- | --- | --- | --- |
| *BoHV-1* | LRS ONT | direct RNA | Moldován et al., 2020 | ENA: PRJEB33511 |
|  | LRS ONT | direct cDNA |  |  |
|  | LRS ONT | amplified cDNA |  |  |
|  | Synthetic LRS Illumina | LoopSeq |  |  |
|  | LRS ONT | direct cDNA | Tombácz et al., 2022b | ENA: PRJEB33511 |
| *EBV* | LRS PacBio RSII | amplified cDNA | O'Grady et al., 2016 | GSE: GSE79337 |
|  | SRS Illumina | amplified cDNA |  |  |
|  |  | CAGE |  |  |
|  | LRS ONT | direct cDNA | Fülöp et al., 2022 | ENA: PRJEB38992 |
|  |  | amplified cDNA |  |  |
| *HCMV* | LRS PacBio RSII | amplified cDNA | Balázs et al., 2017 | ENA: PRJEB22072 |
|  | LRS PacBio Sequel | amplified cDNA | Balázs et al., 2018 | ENA: PRJEB25680 |
|  | LRS ONT | amplified cDNA |  |  |
|  | LRS ONT | direct RNA |  |  |
|  | LRS ONT | CAP-selected | Kakuk et al., 2021 | ENA: PRJEB25680 |
| *HSV-1* | SRS Illumina | amplified cDNA | Rutkowski et al., 2015 | GEO: GSE59717 |
|  |  | amplified cDNA | Pheasant et al., 2018 | SRA: PRJNA505045 |
|  |  | amplified cDNA | Tang et al., 2019 | SRA: PRJNA482043, PRJNA483305, and PRJNA533478 |
|  |  | amplified cDNA | Whisnant et al., 2020 | GEO: GSE128324 |
|  | LRS PacBio Sequel | amplified cDNA | Boldogkői et al., 2018 | ENA: PRJEB25433 |
|  | LRS ONT | amplified cDNA |  |  |
|  |  | direct RNA |  |  |
|  | LRS ONT | direct RNA | Depledge et al., 2019 | ENA: PRJEB27861 |
|  | LRS PacBio RSII | amplified cDNA | Tombácz et al., 2018 | GEO: GSE97785 |
| *PRV* | SRS Illumina | amplified cDNA | Oláh et al., 2015; | ENA: PRJEB9526 |
|  | LRS PacBio RSII | direct cDNA | Tombácz et al., 2016 | ENA: PRJEB12867 |
|  |  | amplified cDNA |  |  |
|  | LRS ONT | amplified cDNA | Moldován et al., 2018 | SRA: PRJNA417577 |
|  | LRS PacBio Sequel | amplified cDNA | Tombácz et al., 2018a | ENA: PRJEB24593 |
|  | LRS ONT | direct RNA |  |  |
|  | LRS ONT | amplified cDNA |  |  |
|  | LRS ONT | amplified Cap-selected |  |  |
|  | LRS ONT | Terminator-handled amplified cDNA | Torma et al., 2021 | ENA: ERP106430 and ERP019579 |
|  | LRS ONT | direct cDNA |  |  |
|  | LRS ONT | direct RNA |  |  |
| *SVV* | LRS ONT | direct RNA | Braspenning et al., 2021 | ENA: PRJEB42868 |
|  | SRS Illumina | amplified cDNA |  |  |
| *VZV* | LRS ONT | amplified cDNA | Prazsák et al., 2018 | ENA: PRJEB25401 |
|  |  | Targeted |  |  |
|  | LRS ONT | amplified Cap-selected | Tombácz et al., 2018b | ENA: PRJEB25401 |
|  | SRS Illumina | amplified cDNA | Braspenning et al., 2020 | ENA: PRJEB38829. |
|  | SRS Illumina | CAGE |  |  |
|  | LRS ONT | direct RNA |  |  |

In the first part of this study, we annotated the exact genomic locations of the canonical viral RNA transcripts and their TIs, including TSSs, TESs, and splice site variants. Previous annotations were either confirmed or in several cases modified on the basis of integration and re-evaluation of the sequencing datasets. We also identified cis-regulatory elements for many of the examined viral RNA transcripts. Intriguingly, promoter elements (TATA boxes) within the Ori sequences were detected in all cases. In this work, we also tried to get a close-to-complete picture about the complexity of TOs at the examined genomic regions. We applied more stringent criteria for transcript annotation at viruses where no dRNA-Seq and/or no CAGE data were available, and therefore we present a lower transcript complexity in these viruses. Relative transcript abundances were determined in viruses for which sufficient data were available. Furthermore, a multi-timepoint real-time RT-PCR (RT^2^-PCR) analysis was applied for monitoring the expression kinetics of three lncRNAs in PRV.

We also applied RT^2^-PCR for the validation of lncRNAs and longer mRNA isoforms of PRV, BoHV-1, EHV-1, HSV-1, and KSHV. Native RNA sequencing was used to validate the results of cDNA sequencing. CAGE-Seq was applied for the validation of KSHV and EHV-1 TSSs obtained by ONT sequencing. We note that the size-biasing effect of the library preparation and the LRS techniques may lead to non-detection or considerable underestimation of the abundance of long (>5kb) transcripts. However, our RT^2^-PCR analysis demonstrated that many of these RNA transcripts are indeed expressed at a low level. The names of orthologous genes differ among the αHVs, therefore, for better comparability, we generally use the names applied in HSV-1 terminology. **Supplementary Table 1** shows the correspondence of orthologous genes.

### 2. Transcripts overlapping or mapping near the replication origins

The polyadenylated transcripts overlapping or mapping proximal to the Oris include lncRNAs, as well as long 5′ and 3′ UTR isoforms of mRNAs (**Figures 2-4, Supplementary Table 2**).

**Figure 2.**
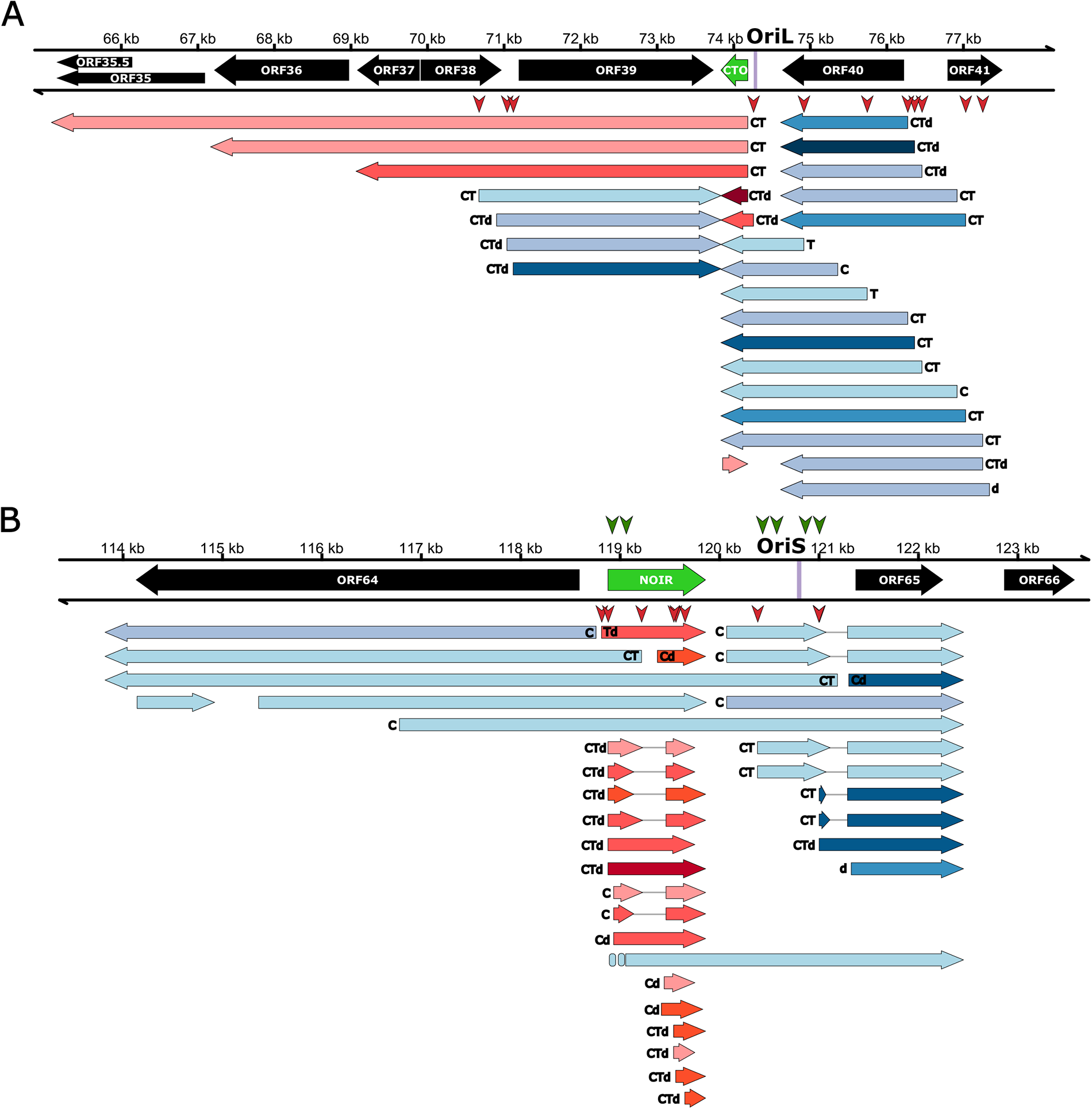
Ori-proximal transcripts of EHV-1. This picture shows the transcripts encoded by the OriL-(A) and OriS-proximal regions of Equid herpesvirus 1. All the putative transcripts were identified by LoRTIA software using dcDNA datasets, unless indicated otherwise. Protein coding genes are labeled with black arrows, non-coding genes with green arrows, mRNAs with blue arrows, and ncRNAs with red arrows. The relative transcript abundance was labeled by shades. We carried out CAGE-Seq (transcript detected by this technique is labeled by ‘C’). Those transcripts that have a proximal TATA box are marked by a ‘T’ letter, and the vertical red arrows indicate the positions of TATA boxes on the genome. Those transcripts that were also detected by dRNA-Seq are marked with a ‘d’ letter at the upstream positions. Introns are indicated by horizontal lines.

**Figure 3.**
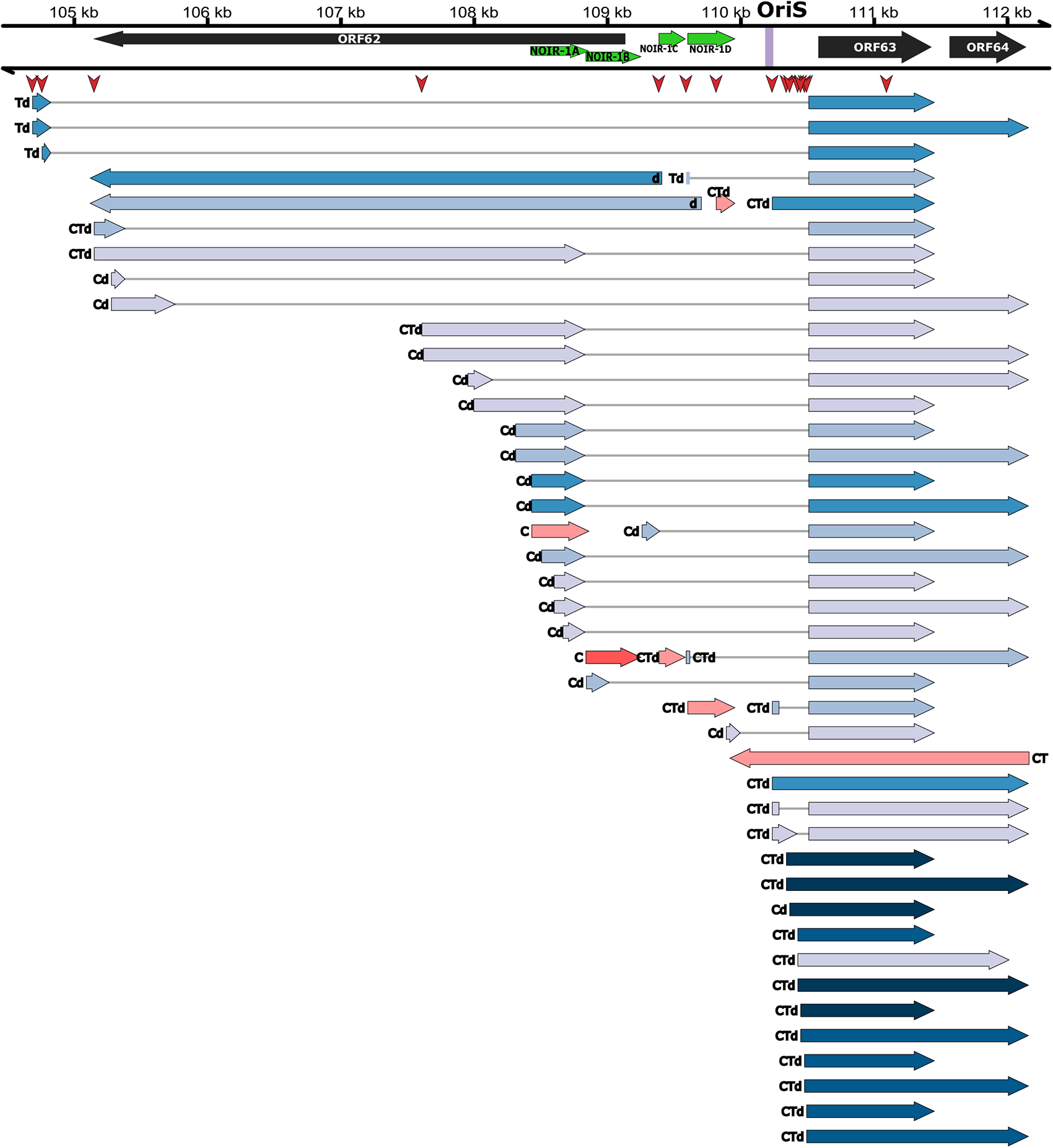
OriL-proximal transcripts of VZV. This picture shows the transcripts encoded by the OriS-proximal region of Varicella-zoster virus. All of the putative transcripts were identified by LoRTIA software using dcDNA datasets, unless indicated otherwise. Protein coding genes are labeled with black arrows, non-coding genes with green arrows, mRNAs with blue arrows, and ncRNAs with red arrows. The relative transcript abundance was labeled by shades. Transcripts that have a proximal TATA box are marked by a ‘T’ letter, and those that were also detected by dRNA-Seq are marked with a ‘d’ letter at the upstream positions. The vertical red arrows indicate the positions of TATA boxes on the genome. Transcripts that were detected by CAGE-Seq are marked by a ‘C’ letter. Introns are indicated by horizontal lines.

**Figure 4.**
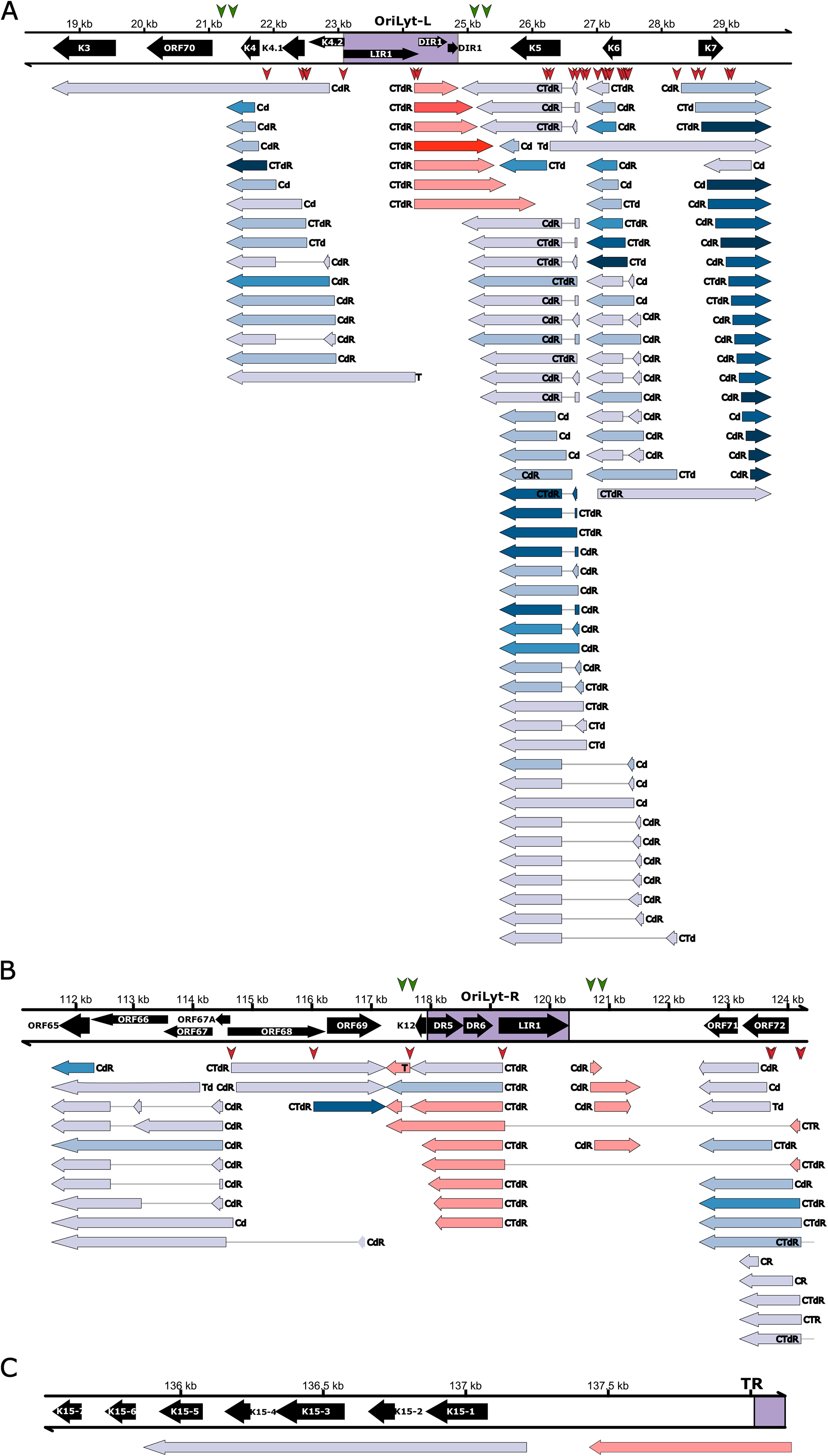
Ori-proximal transcripts of KSHV. This picture shows the transcripts specified by the Ori-proximal regions of Kaposi’s sarcoma-associated herpesvirus (Orilyt-L: A; OriLyt-R: B; TR: C). All of the putative transcripts were identified by LoRTIA software using dcDNA datasets, unless indicated otherwise. Protein coding genes are labeled with black arrows, non-coding genes with green arrows, mRNAs with blue arrows, and ncRNAs with red arrows. The relative transcript abundance was labeled by shades. Transcripts that have a proximal TATA box are marked by a ‘T’ letter, and those that were also detected by dRNA-Seq are marked with a ‘d’ letter at the upstream positions. The vertical red arrows indicate the positions of TATA boxes on the genome. Transcripts that were detected by CAGE-Seq are marked by a ‘C’ letter. RAMPAGE data were also available for KSHV (transcripts detected by this technique are marked by an ‘R’ letter). Introns are indicated by horizontal lines.

#### Alphaherpesviruses – OriS

In this part of the study, we identified novel transcripts and TIs in the vicinity of the OriSs of α-HVs. **Figures 2 and 3, Supplementary Figures 2-5, and Supplementary Table 2** show the complete list of the annotated transcripts, including lncRNAs, such as the OriS-RNA of BoHV-1, the OriS-RNA1 of HSV-1, the NOIR-1 transcripts (in PRV, EHV-1, VZV and SVV), and NOIR-2 transcript (in PRV). We also found that the long 5′-UTR TI of the *us1* gene (in all of the examined αHVs) and of the *icp4* gene (in BoHV-1, EHV-1, HSV-1 and SVV) overlaps the OriS. We cannot exclude that ICP4 TIs in other αHVs also overlap the OriS, but they might have gone undetected due to their long size and low abundance. In HSV-1 TRS region, the 5′ UTR region of the US12 long TI also forms an overlap with the OriS. We detected a very complex splicing pattern of US1 transcripts in αHVs. Novel lncRNAs oriented antisense to the HSV-1 OriS-RNA1 were also identified.

Consensus promoter elements (TATA boxes) for many of these transcripts were identified. Members of the NOIR-1 family are arranged in divergent configuration with respect to the *icp4* gene. The canonical version of these RNAs does not overlap *icp4* whereas the longer NOIR-1 variants form partial overlap with this major TR gene. In EHV-1, VZV, SVV, and probably in PRV, a long TI of US1 is initiated from the promoter of *noir* gene. In SVV, we identified a long TSS variant of NOIR-1, which partially overlaps the canonical ICP4 transcript and completely overlaps the long TSS isoform of ICP4 isoform. Both this and the canonical NOIR-1 overlap the OriS. While NOIR-1 is expressed at a moderate level, NOIR-2 positioned in convergent orientation with respect to NOIR-1 is expressed in a very low-abundance. NOIR-2 has only been detected in PRV. In VZV, we detected five lncRNAs with distinct TSSs and TESs at this genomic region. We termed them as NOIR-1A, -1B, 1C, -1D, and -1E. We note here that VZV NOIR-1 transcripts were confirmed with lower level of evidence than other transcripts. We identified TIs with TSSs mapping very close to the TATA boxes located within the OriSs in all of the six αHVs, which indicates that these promoter elements are functional.

##### a. Alphaherpesviruses – OriL

The CTO transcript family has been described by Tombácz and colleagues^15^. In the original publication, three 3′-coterminal transcripts have been identified, the short CTO-S, the CTO-M, which is initiated near the poly(A) signal of *ul21* gene, and CTO-L, which is a transcriptional read-through TI (3′ UTR variant) encoded by the *ul21* gene. We note that the long 3′ UTR isoforms with unique TES are extremely rare in αHV mRNAs. The list of this family has been updated later^58^ and a more detailed update is published in this current report. CTO transcripts were also detected in EHV-1 (**Figure 3**; **Supplementary Figures 3 and 5, and Supplementary Table 2**), but not in other herpesviruses with annotated transcriptome. CTO-S is expressed in a very high abundance in both PRV and EHV-1. We also discovered antisense transcripts (asCTO) that overlap the CTOs in both viruses. Additionally, we detected a tail-to-tail (convergent) TO between the 3′ UTR isoforms of CTO-S and UL22 transcripts, and very long read-through CTO transcripts were also described in both viruses. We analyzed two PRV strains, the laboratory strain Kaplan (PRV-Ka^68^) and a field isolate (strain MdBio: PRV-MdBio^69^). In HSV-1, both members of the divergent *ul29*-*ul30* gene pair generate long 5′ UTR variants that overlap the OriL. No lncRNA was detected near the HSV-1 OriL.

##### b. Betaherpesviruses

**Supplementary Figure 6 and Supplementary Table 2** show the HCMV transcripts around the OriLyt. Our data show that one of the major lncRNAs, RNA4.9^46^ is initiated from the OriLyt. The ul59^70^, SRT^49^ and one of the two vRNAs (vRNA-2)^50^ were also identified using our previously published dataset^37^. In our work, we were unable to detect the SRT and the vRNAs using LRS, probably because of their small sizes. However, we detected two longer versions of UL58 lncRNA and a shorter isoform of UL59 lncRNA.

##### c. Gammaherpesviruses

**Figure 4, Supplementary Figure 7, and Supplementary Table 2** show the list of EBV transcripts around the three Oris. It can be seen in **Panel A of Supplementary Figure 7 t**hat the long 5′ UTR isoform of *BCRF1* gene overlaps OriP, the latent replication origin of the virus^71^. Likewise, the long 5′ UTR variants of *BHRF1* gene overlap OriLyt-L. One of these transcripts is also a splice isoform encoded by this gene. The promoter of *BHLF1* gene is located within the Orilyt-L^51^. Here we describe novel isoforms of lncRNAs that either overlap the OriLyt-R with their introns or are initiated from the replication origin. Our analyses revealed that several OriLyt-L-associated ncRNAs of varying length can be produced from the same TSS besides the 1.4-kb ncRNA during lytic reactivation of KSHV in primary effusion lymphoma cells **Figure 4, Supplementary Table 2).** Whether they all have the same role in viral DNA replication remains to be tested. The OriLyt-L is flanked by short protein-coding genes such as K4.2, K4.1 and K4 on the left and K5, K6, and K7 on the right side. These genes have a wide range of functions in the regulation of lytic replication, cytokine signaling, protein degradation, cell cycle and apoptosis^72^. Previous studies indicated that K4, K4.1, and K4.2 are transcribed as mono-, bi-, and tricistronic mRNAs^73^, but our analysis show a more complex expression pattern that includes unspliced RNAs of variable length and spliced RNA variants. We also found that K5/K6 genes can be expressed not only individually but also through splicing, which results in mRNAs with a first exon of different length. Importantly, our results are in line with previous genomics studies but also expand on the number of different viral RNA transcripts that can be produced from the OriLyt-L locus, which can potentially increase the coding capacity of viral mRNAs^74–76^. KSHV latency locus is located between K12 and LANA (ORF73), which encodes 4 protein-coding latent genes (K12, K13, ORF72, ORF73) and 12 pre-miRNAs. This locus also includes OriLyt-R, which is not involved in lytic viral DNA replication. Previous studies have demonstrated that the latent genes are products of a complex gene transcription, which also includes alternative splicing^76–78^. The viral miRNAs are encoded in an intron, which is part of viral RNAs that are initiated from a promoter upstream of ORF72. While we confirmed that this splicing takes place, we also detected several ncRNAs that are antisense with the miRNA-coding genomic regions. It will be interesting to test in future studies whether any of these ncRNAs can interact with the viral miRNAs to regulate their expression and function.

### 3. Overlapping RNA isoforms of transcription regulator genes

The *us1* gene encodes very long 5′ UTR isoforms, which were found to form head-to-head TOs with the ICP4 transcripts in HSV-1, EHV-1 and VZV (**Figure 2**), but TOs may also be formed in other αHVs. In HSV-1, the 5′ UTR isoforms of US10-12 polycistronic transcripts were found to form a divergent TO with the *icp4* gene. In HSV-1, we also detected an ICP4 TIs with long 5′ UTR, which overlaps the *us1* gene. Additionally, a 3′ UTR isoform of *icp4* has been described to form a parallel TO with the downstream *icp0* gene in BoHV-1, however, the *icp4* ORF is spliced out from this transcript resulting a chimeric transcript containing the full-length *icp0* gene and part of the 5′ UTR of ICP4 transcript^79^. Additionally, ICP4 TIs were detected to form similar chimeric transcripts and also bicistronic transcripts with the BoHV-1 CIRC RNA mapping to the adjacent genomic locus in the circular or concatemeric viral genome. It is possible that these transcripts form an overlap with the convergent UL54 transcript, but in this study, we obtained no evidence for this.

### 4. Long non-coding RNAs mapping within or proximal to the transcription regulator genes

Besides the long mRNA isoforms encoded by the TR genes, lncRNAs with a putative regulatory function have also been described at these genomic loci. Antisense RNAs (asRNAs) overlapping the *us1* gene (termed asUS1) have been detected in BoHV-1 and PRV. ELIE has previously been detected only in PRV, but we could identify a transcript with a similar genomic location in EHV-1 (**Figure 2**). The lncRNA ELIE is mapped between the *icp4* and *ep0* genes. One of its TIs is 5′-coterminal with the NOIR-1 transcripts. We also detected an asRNA in EHV-1, designated as as64, with the same orientation as PRV ELIE, but within the *icp4* gene. May be these two transcripts have a similar function. A HSV-1 transcript initiated at the 3′ end of *icp4* gene and terminated at the us1 gene is also described in this study. Furthermore, we detected a TSS of long 5′ UTR isoform of VZV *us1* gene, which is mapped to a downstream position of *icp4* gene, that is, at the homologous position as the TSS of PRV ELIE. AZURE, another lncRNA in PRV stands at opposite direction to the *us1* gene. The LAT, LLT, and L/ST lncRNAs are expressed during latency^80^.

### 5. Transcriptional overlaps of replication genes

In Simplexviruses, the long 5′ UTR isoforms of divergent *ul29*-*ul30* gene pair overlap not only the OriL but also each other (**Figure 3**). Intriguingly, both genes code for regulators of DNA replication: the *ul29* specifies the major DBP whereas the *ul30* encodes a DNP. Thus, it seems that these genes act not only through producing replication regulator proteins but also their transcripts and transcriptions potentially interfere with each other and also with the replication initiation at the OriL. In both HCMV (*ul57*) and human herpesvirus type 6 (HHV-6) (*ul42*), the *ul29* orthologs are adjacent to the OriLyt.

Intriguingly, in αHVs, three ‘hard’ TOs between gene pairs are present, and one of the partners in these is always a gene playing a role in viral replication. These TOs are as follows: *ul30*/*ul31*, *ul6*-*7*/*ul8-9*, *ul50*/*ul51* (*ul30*: DNP; *ul8*: DNA helicase; *ul9*: OBP; *ul50*: deoxyuridine triphosphatase). It is assumed that the transcriptional interference between gene partners with ‘hard’ convergent TOs is stronger than those with ‘soft’ TOs.

### 6. Defining the TSS patterns of the examined genomic regions

Determining the TSS of RNA molecules is a major challenge of transcriptome research. Besides the application of various LRS and SRS approaches, we also performed CAGE sequencing in the case of EHV-1 and KSHV (**Figure 5**) and used other’s data for VZV for this purpose^81^. TSSs detected by CAGE-Seq are also indicated in **Figures 2-4**. Our CAGE-Seq results on the TSSs of KSHV were compared with the results obtained by RAMPAGE-Seq published by others ^82^. We described altogether 199 KSHV transcripts of which 192 were confirmed by CAGE and 159 by RAMPAGE. All of the TSSs detected by RAMPAGE were also identified by CAGE.

**Figure 5.**
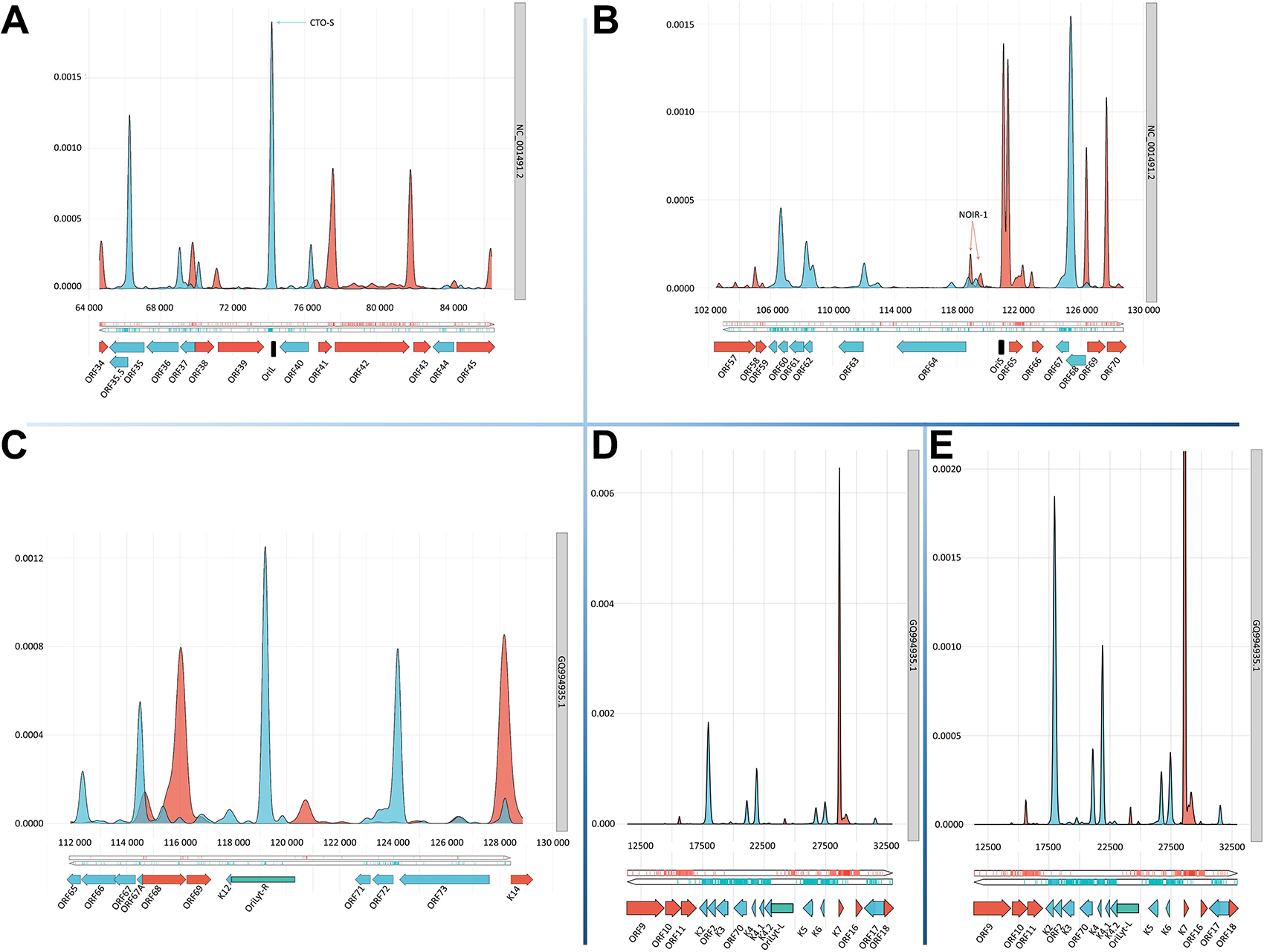
Distribution of TSSs at the examined genomic regions determined by CAGE-Seq. TSS distributions are illustrated in the following genomic regions of EHV-1 and KSHV: **A.** EHV-1 OriL; **B.** EHV-1 OriS of the IRS; C. KSHV OriLyt-R; **D.** KSHV OriLyt-L; **E.** KSHV OriLyt-L – here a higher resolution is used for the better visibility of the low-abundance TSSs. It can be seen that CTO-S transcript is at a high level (A), whereas NOIR-1 is a group of transcripts expressed in a relative low level. Smoothed density plots of the 5′ ends in the CAGE data. Here, y-axis shows the result of the probability estimation of the 5′ ends using a probability density function (details are described in the Materials and Methods section). Coding sequence (CDS) annotations for the respective genomes (shown with the accession number at the right) were visualized in the lower part. The coverage of the positive strand and the CDS annotation are shown in red while these were illustrated in blue in the negative strand. The Ori regions in EHV-1 are illustrated in black whereas in KSHV they are depicted in green – in the latter case, an accompanying white box shows the 20-nt binding site for the DNA replication origin-binding protein.

Our study revealed that while the strong promoters produce small, the weak promoters produce high TSS variance.

### 7. Transcript validation using RT^2^-PCR

The most important novel transcripts of five viruses (PRV, BoHV-1, EHV-1, HSV-1, and KSHV) were further validated using RT^2^-PCR (**Figure 6**). The Ct values clearly indicate that the long TSS isoforms of the TR genes overlapping the Ori and each other are expressed at a low-level.

**Figure 6.**
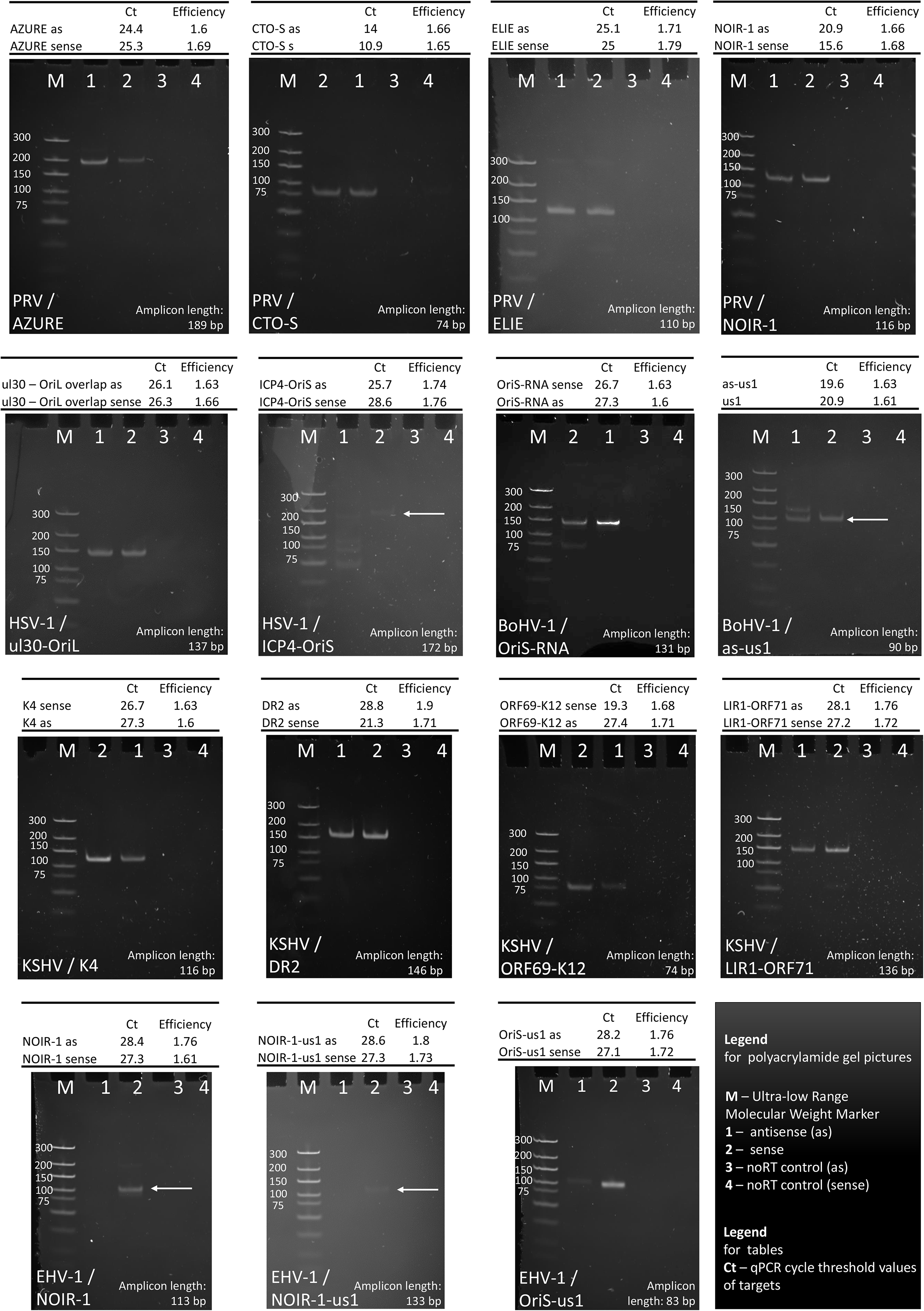
Validation of Ori-adjacent transcripts using RT^2^-PCR. In this figure, we used RT^2^-PCR and gel electrophoresis for the validation of important Ori-adjacent transcripts. The name of the virus and the transcripts are indicated at the appropriate panels. We examined the transcriptional activity of both DNA strands, and used no RT controls for each transcript. The lanes are as follows in every panel. M: molecular weight marker; 1: antisense transcripts; 2: sense transcripts (mRNA, or the canonical lncRNA); 3: No RT for the antisense transcripts; 4. No RT for the sense transcripts. We also indicated the amplicon lengths as well as the Ct and efficiency values, which allow the estimation of the quantity of the transcripts.

### 8. Transcription kinetics of lncRNAs

In this work, we carried out kinetic analyses using three PRV lncRNAs, which are as follows: CTO-S, NOIR-1 and AZURE (**Figure 7**). We obtained that CTO-S (R_4h/2h_ = 10.6; R_8h/4h_ = 17.4) and NOIR-1 (R_4h/2h_ = 11.9; R_8h/4h_ = 12.7) are expressed at a low level at the first four hours of post-infection while AZURE (R_4h/2h_ = 27.3; R_8h/4h_ = 5.2) is expressed at high level at 4h of post-infection. According to these results, CTO-S and NOIR-1 are categorized as late genes whereas AZURE as an early non-coding gene.

**Figure 7.**
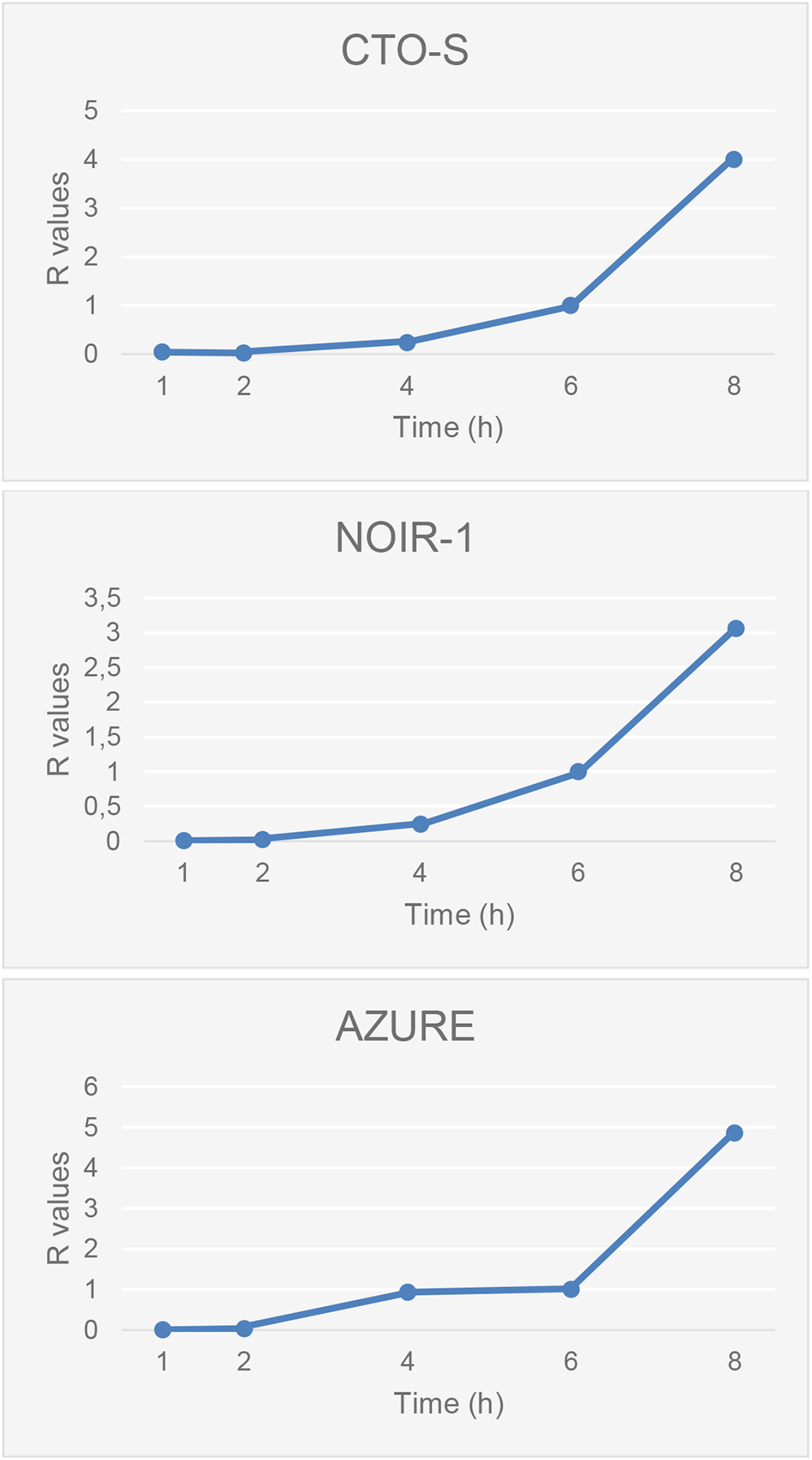
Expression kinetics of three lncRNAs of PRV. This figure shows the transcription kinetics of three PRV lncRNAs, which are the CTO-S, NOIR-1, and AZURE. The CTO-S and NOIR-1 have a late while the AZURE has an early expression kinetics.

## DISCUSSION

With the emergence of LRS technologies, it is now feasible to identify and precisely annotate transcripts and RNA isoforms, including length and splice variants. Single Molecule, Real-Time (SMRT) sequencing with PacBio^62, 83^, nanopore sequencing with ONT^16^, and LoopSeq synthetic LRS (running on Illumina platform) with Loop Genomics^56^ have already been used alone, or in combination with each other, or with SRS for viral transcript annotation in all three subfamilies of herpesviruses (HSV-1^84, 85^; VZV^81, 86^; PRV^15^; BoHV-1^87^; HCMV^61^; EBV^62, 63^).

In recent studies, we and others have demonstrated the existence of a previously hidden, highly complex, genome-wide network of transcriptional overlaps in different viral families^15, 22, 62, 85, 88, 89^. It has been shown that the RNA molecules encoded by closely spaced genes overlap each other in parallel, divergent, and convergent manners. This phenomenon implies an interaction between the transcription machineries at the TOs throughout the entire viral genome^90^. We and others have formerly demonstrated that the replication origins of several viruses form overlaps with specific lncRNAs and with long 5′ or 3′ UTR isoforms of mRNAs^41^. We note that many of the 5′ UTR isoforms are possibly non-coding due to the long distance between the TSS and start codon. Exceptions may be those transcripts whose a large part of the 5′ UTR are spliced out. Functional analyses have revealed the mechanistic details of how replication RNAs control the onset of DNA synthesis through the formation of RNA:DNA hybrids in several viruses^47^.

In this work, we present the state-of-the-art of our current knowledge focusing on the structural aspects of Ori-overlapping and Ori-adjacent transcripts and transcript isoforms that potentially play key roles in the regulation of replication and genome-wide viral transcription of herpesviruses. We put a more emphasis on the examination of αHVs because less information is available for such transcripts in these viruses. Using novel and previously published sequencing data, we discovered new transcripts and corrected the earlier annotations of the already described RNAs. We also identified an intricate meshwork of TOs between transcripts encoded by replication and transcription regulatory genes and by specific lncRNAs around them. We detected promoter consensus elements within the Oris in all of the examined herpesviruses. Although the Oris of herpesviruses contain AT-rich sequences, which can be misidentified as TATA boxes, we detected the corresponding transcripts with a TSS being proximal to these cis-regulatory sequences in all cases. CAGE analyses also indicated that these sequences function as promoters. The terminology of the reported lncRNAs is somewhat arbitrary as, unlike protein-coding genes, their sequences and locations are poorly conserved due to mapping to the rapidly evolving intergenic regions. Furthermore, it is possible that some of these lncRNAs with the same name (e.g. NOIR-1) have a polyphyletic origin. However, the CTO-S and NOIR-1 are apparently orthologous in PRV and EHV-1.

The co-temporal activity of DNA replication and transcription at the same genomic region generates interference between the two machineries^91^. Data suggest that the collisions lead to more dramatic consequences if occurring in convergent orientation than co-directionally^92^. Several molecular mechanisms have evolved to minimize the conflict between the RNP and the replication fork^93^. Our current study, however suggests the existence of a novel intrinsic regulatory system based on an interaction between the two apparatuses for controlling the initiation of both replication and transcription in a cooperative manner. This interaction is supposed to be mediated through the clash and competition between the transcribing RNP and the replisome, and through the inhibition of the assembly of pre-replication and transcription initiation complexes by the ongoing processes of replication and transcription (**Figure 8** **and Supplementary Figures 8-9**).

**Figure 8.**
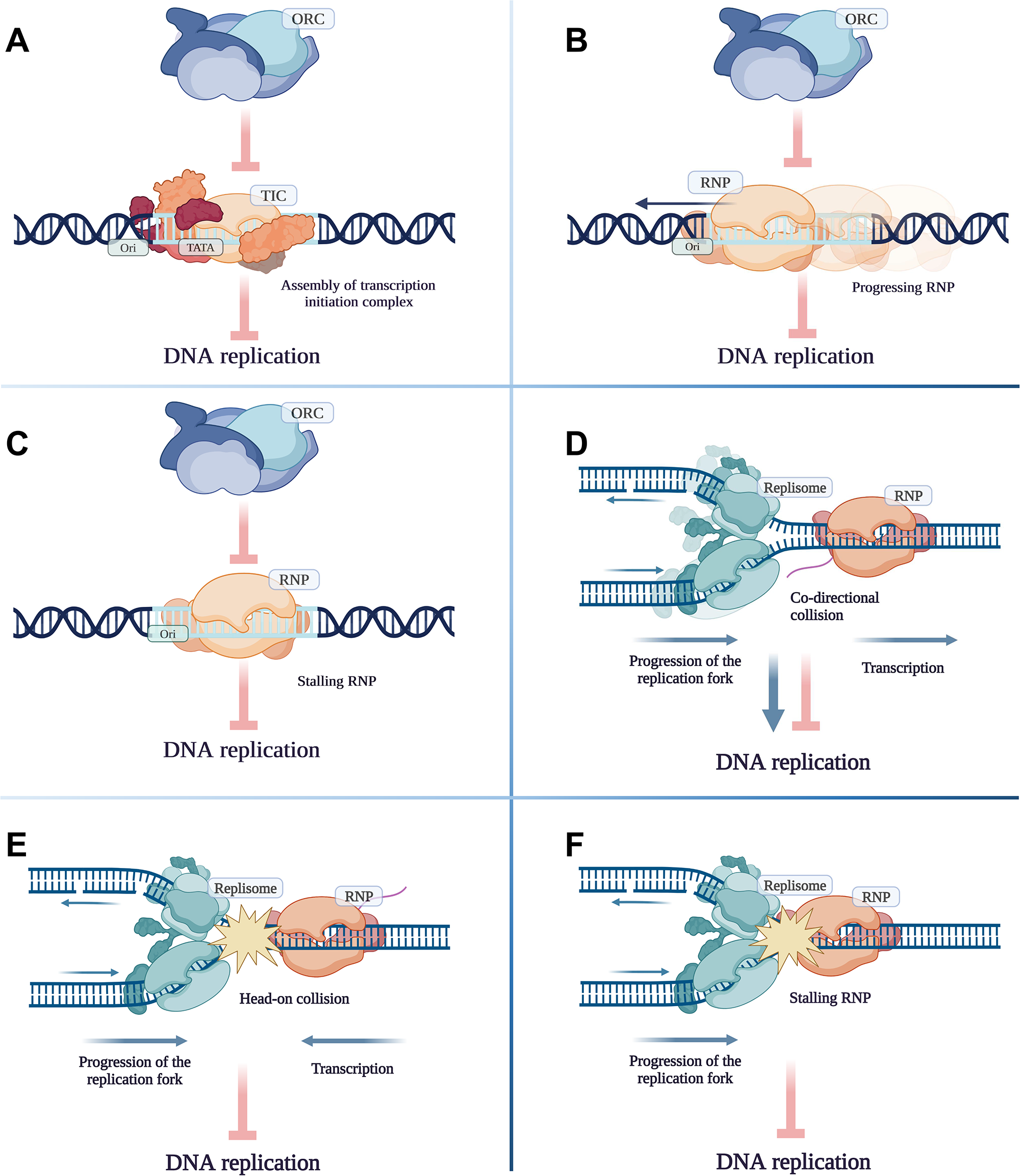
Potential effects of transcription on the DNA replication. A. The assembly of transcription initiation complex on promoters located within the replication origin inhibits the assembly of ORC and replisome. B. Passage of RNA polymerase II across the replication origin inhibits the assembly of ORC and replisome. C. Stalling of RNA polymerase II on the replication origin inhibits the assembly of ORC and replisome. D. Co-directional movement of replication and transcription machineries can slow down or speed up the progression of replication fork. Transcription can facilitate DNA replication by pre-opening the two DNA strands. E. Head-on collision of replication and transcription machineries inhibits both processes. F. Stalling of RNA polymerase II inhibits the progression of replication fork.

In αHVs, the position of OriS is highly conserved on the genome: it is located in the US repeats and surrounded by the major TR genes, *icp4* and *us1* (except in Simplexviruses, where *us1 is* substituted by *us12* in the terminal IR). Here, we demonstrated that the TR genes encode transcript isoforms which overlap not only the OriS but also the transcripts of the opposite TR gene and/or specific proximal lncRNAs. The probable function of the TOs is to allow additional forms of interplay between these genes – besides the TF/promoter interaction – which is supposed to be achieved by RNA:DNA hybridization, and possibly by RNA:RNA hybridization, as well as by the interference between the transcription apparatuses. Thus, the US IR region of αHVs appears to function as a ‘super regulatory center’, where the initiation of both DNA replication and global transcription is collectively controlled through a mutually interacting multilevel system. This region is far the most complex genomic locus of αHVs, which encodes intergenic lncRNAs, including intergenic transcripts and asRNAs, long TIs of mRNAs, and an intricate TO pattern of the local transcripts. Further, besides the lytic transcripts, several latent lncRNAs (LAT, LLT, L/ST) and miRNAs are also expressed from this genomic region^94^.

The NOIR-1 transcript family mapped to the US IR is the evolutionary innovation of the Varicellovirus genus. The exact location of these lncRNA molecules exhibit a significant variance. In PRV, the long TIs of NOIR-1 overlap the *icp4*, but no OriS overlapping TES TI has been detected in this virus. The probable reason for this is that NOIR-1 is opposed to NOIR-2, a unique in PRV RNA, which might prevent the read-through of NOIR-1. The PRV NOIR-1 is 3′ co-terminal with the LLT transcripts that are expressed during latency. In EHV-1, the NOIR-1 long TI overlaps the *icp4* while the long TSS isoform of US1 transcript, utilizing the NOIR-1 promoter, overlaps the OriS. The VZV NOIR-X promoters also control the transcription of the long TSS variants of US1, however, the *icp4* overlapping is provided by NOIR-1. SVV has evolved another mechanism: NOIR-1 overlaps the *icp4* and alongside with the short TES isoforms of NOIR-X, it is terminated at the OriS. Similar to VZV, the TSS isoform of US1 RNAs utilizes the NOIR-1 promoter SVV and overlaps the OriS. BoHV-1 lacks NOIR-1, but it has a long TSS isoform of US1 transcripts that overlaps the OriS. Additionally, BoHV-1 expresses OriS-RNA whose orientation is parallel to the *icp4*, and its promoter is located within the OriS. In HSV-1, the OriS-RNA1 has an opposite polarity than the NOIR-1 and is located on the other side of the OriS. The promoter of this transcript also controls the long TSS isoform of the ICP4 transcript. The NOIR-1 transcripts may be involved in the control of transcription through overlapping the *icp4* gene, and/or in the control of DNA replication through overlapping the OriS (in the case when they are the upstream part of the US1 TI), or in the control of both transcription and replication since some of them overlap both OriS and *icp4*. The same might also be true for the HSV-1 OriS-RNA1 because it also serves as the 5′ UTR part of the long ICP4 TI.

PRV has a low-abundance lncRNA, termed ELIE whose long TES isoform is co-terminal with NOIR-1. Another lncRNA, the EHV-1 as64 overlapping the 3′ region of *icp4*, may be the functional homologue of PRV ELIE. Furthermore, we detected asUS1 transcripts that overlap the *us1* gene partially in PRV and fully in BoHV-1. The PRV AZURE asRNA overlaps the *us3* and *us4* genes. Although we could not detect transcriptional read-through from the *us3/us4* genes toward the *us1* gene and *vice versa*, it is known that eukaryotic transcription proceeds through the TES, which is followed by the removal of the read-through part of the RNA molecule.

While the OriS is essential in αHVs, the OriL appears to be dispensable since some Varicelloviruses, e.g. VZV and BoHV-1, do not have it. In addition, OriL has been relocated in PRV and EHV-1 compared to Simplexviruses. The original genomic position of OriL is obviously the intergenic region between two main replication genes (*ul30* and *ul29*), because in the distantly-related βHVs (HCMV and HHV-6) the OriLyt is adjacent to the *ul29* ortholog.

It may not be a coincidence that the OriL of Simplexviruses is surrounded by the main replication genes. We hypothesize that these genes might interact with the replication not only through the conventional TF/promoter binding but also at other levels including the RNA:DNA and/or RNA:RNA hybridization, and also through the interference of the transcription and replication machineries at the overlapping region.

CTO transcripts have been previously identified in PRV^54^. Here, we report the detection of the orthologous transcripts in EHV-1. The canonical CTO-S does not overlap the OriL, but it might interfere with the replication by helping to separate the two DNA strands and thereby also determining the orientation of replication. One of the long CTO-S TIs is controlled by a promoter within the OriL and another one is the TES variant of *ul21* gene.

The OriLyt-L of EBV is adjacent to the non-coding BHLF1 gene, which is a latency regulator^95^ while the OriLyt-R is adjacent to LF2, which is an inhibitor of replication^96^.

The OriLyt-L of CMV is overlapped by lncRNAs among which the RNA4.9 has been shown to regulate DNA replication^47^. We have described novel Ori-overlapping lncRNAs of HCMV previously^37^ and also in this recent report.

The KSHV LANA plays an essential role as a latency regulator^97^. In this study, we describe additional Ori-overlapping transcripts, including lncRNAs and long 5′ UTR isoforms of mono- and polycistronic mRNAs.

We also observed a potentially intriguing phenomenon: in αHVs, altogether three ‘hard’ convergent TOs are formed, which supposedly represent a strong confrontation between the transcription apparatuses during the synthesis of RNA molecules. We noticed that one of the partners forming these TOs is always an auxiliary replication gene. The involvement of such genes in every ‘hard’ TOs may not only be a mere co-occurrence but may also have a biological relevance, which is currently unknown. However, we can speculate that the ‘hard’ TO between the *ul30**/**ul31* partner genes of HSV-1 allows *ul31* gene to generate an OriL-overlapping long TES isoform through transcriptional read-through. Indeed, we detected a high-frequency transcriptional read-through in this gene using RT^2^-PCR. Intriguingly, although in PRV and EHV-1 the OriL is located between the *ul21* and *ul22* genes, the OriL is also overlapped by a long TES isoform (of *ul21*). Since no long TES isoforms of *ul21* in Simplexviruses and of *ul31* in Varicelloviruses are produced, it is reasonable to assume that these types of TI were created for the interference with the initiation of DNA replication.

The amount of particularly long 5′ UTR variants was quantified using RT^2^-PCR and we found that their low abundance cannot be explained by the size-bias of the sequencing alone, but these transcripts are indeed expressed at a low level. Although many of these transcripts comprise complete ORFs, they are probably ncRNAs since their transcription and translation start sites are far apart from each other. Hence, they might solely be transcriptional by-products without functioning as protein coding sequences, or even as RNA molecules. In other words, their transcription may be important by regulating the transcription of other RNA molecules, or the DNA replication.

In conclusion, herpesviruses, even the closely related species, evolved distinct solutions for the formation of TOs at the Oris and also the TR genes, which indicates the importance of this phenomenon in the control of DNA replication and global transcription. Whilst at the OriS region of αHVs, the major TR genes appear to control each other and the initiation of DNA replication, at the OriL region of Simplexviruses, the main replication genes might regulate each other and also the replication through mechanisms acting on TOs. These putative mechanisms provide multiple regulatory layers in addition to the conventional transcription factor/promoter interaction. The ICP4 TR has been shown to facilitate the expression of *icp0*^98^ genes by binding to its promoter. The ICP22 (*us1* product) inhibits the expression of both *icp4* and *icp0*^99^. Furthermore, ICP0 converts ICP4 from a repressor to an activator of mRNA synthesis in HSV-1^100^. The mutation of *us1* gene leads a differential effect on the transcription kinetics of E and L genes^101^.

We believe that the significance of our results and their implications have a much broader perspective and represent a general way of how the regulation of herpesviral replication and transcription co-evolved with each other.

## METHODS

### Cells and Viruses

Besides our novel data, we also used several other datasets for the analyses in this study (**Table 1**).

#### PRV

For the generation of novel transcript data, we used the following three immortalized cells for the propagation of strain MdBio of pseudorabies virus (PRV-MdBio^69^): PK-15 porcine kidney epithelial cell line (ATCC® CCL-33™), C6 cell line derived from a rat glial tumor (ATCC® CCL-107™), and PC-12 (ATCC® CRL-1721™) derived from a pheochromocytoma of the rat adrenal medulla, that have an embryonic origin from the neural crest. Each experiment was carried out in three biological replicate. PK-15 cells were grown in Dulbecco’s modified Eagle medium (DMEM) (Gibco/Thermo Fisher Scientific), supplemented with 5% fetal bovine serum (FBS; Gibco/Thermo Fisher Scientific) and 80 *μ*g of gentamycin per ml (Gibco/Thermo Fisher Scientific) at 37°C in the presence of 5% CO_2_. C6 cells were cultivated in F-12K medium (ATCC), complemented with 2.5% FBS and 15% horse serum (HS; Sigma-Aldrich) at 37°C in the presence of 5% CO_2_. PC-12 cells were maintained in RPMI-1640 medium (ATCC) supplemented with 5% FBS and 10% HS at 37°C in the presence of 5% CO2. Virus stock solution was prepared as follows: PK-15 cells were infected with 0.1 multiplicity of infection [MOI = plaque-forming units (pfu)/cell]. Viral infection was allowed to progress until complete cytopathic effect was observed, which was followed by three successive cycles of freezing and thawing of infected cells in order to release of viruses from the cells. Each cell type was infected with 1 MOI of PRV-MdBio. Infected cells were incubated for 1 h at 37 °C followed by removal of the virus suspension and washing the cells with phosphate-buffered saline (PBS). Following the addition of new culture medium, the cells were incubated for 0, 1, 2, 4, 6, 8, or 12 h. Following the incubation, the culture medium was discarded and the infected cells were frozen at −80°C until further use.

For the kinetic analysis, we used 1 MOI of PRV-Ka for the infection of PK-15 cells. Cells were first incubated at 4 ◦C for one hour for synchronization of infection, and then placed in a 5% CO2 incubator at 37 ◦C. Infected cells were collected at every 30 minutes within an 0 to 8 h interval. Cells were washed with PBS, scraped from the culture plate, and centrifuged at 3000 RPM for 5 min at 4 ◦C.

#### EHV-1

Equid alphaherpesvirus 1 (EHV-1) was also used in this study in order to obtain comprehensive view on global herpesvirus transcriptomes. For this, a field isolate of the virus (EHV-1-MdBio) was used, which is originated from Marócpuszta (Hungary) from the organs of an aborted colt fetus in the 1980s. A confluent rabbit kidney (RK-13) epithelial cell line (ECACC 0021715) was used for viral propagation. The experiments were carried out in three technical replicates.

RK-13 cells were maintained in DMEM (Sigma). The media was supplemented with 10% fecal calf serum (FCS, Gibco). The culture conditions were as follows: 37 °C in the presence of 5% CO_2_. A virus stock solution was prepared by infecting the cells with EHV-1-MdBio at MOI of 0.1. Three freeze-thaw cycles were applied to release the viruses from the cells. For the EHV-1 long-read RNA-seq investigations, RK-13 cells were infected with 4 MOI of the virus. Three technical replicates were carried out. Viral infected RK-15 cells were incubated at 4 °C for 1 h, then the virus suspension was removed and cells were washed with PBS. Next, new media was added to the cells, which were incubated for 1, 2, 4, 6, 8, 12, 18, 24 or 48 h. Finally, the culture medium was removed from the cells and they were stored at -80 °C until further use.

##### KSHV

The KSHV-positive primary effusion lymphoma cell line iBCBL1-3xFLAG-RTA^102^ was maintained in RPMI 1640 medium, which was supplemented with 10% Tet System Approved FBS (TaKaRa), penicillin/streptomycin, and 20 μg/ml hygromycin B. KSHV lytic reactivation was induced by treating 1 million of iBCBL1-3xFLAG-RTA cells with 1 μg/ml doxycycline for 24 h. For measuring KSHV gene expression by RT-qPCR, cDNA was generated with iScript cDNA Synthesis kit (Bio-Rad) followed by SYBR green-based real-time quantitative PCR analysis using gene specific primers. The relative viral gene expression was calculated by the delta-delta Ct method where 18S was used for normalization. The sequences of the primers have been reported previously (Toth et al., 2013). The following antibodies were used for immunoblots: anti-FLAG (Sigma F1804), anti-ORF6 (from Dr. Gary S. Hayward, Johns Hopkins University), and anti-Tubulin (Sigma T5326).

##### VZV

Human primary embryonic lung fibroblast cell line (MRC-5) obtained from the American Type Culture Collection (ATCC) was used for the propagation of the live attenuated OKA/Merck strain of varicella zoster virus (VZV). Cells were grown in DMEM supplemented with antibiotic/antimycotic solution and 10% fetal bovine serum (FBS) at 37 °C in a 5% CO_2_ atmosphere. Infected cells were harvested by trypsinization when they displayed cytopathic effect (after 5 days).

##### HSV-1

An immortalized kidney epithelial cell line (Vero) was used for the propagation of HSV-1. The cell culture was grown in DMEM (Gibco/Thermo Fisher Scientific) with 10% Fetal Bovine Serum (Gibco/Thermo Fisher Scientific) and 100 μl/ml penicillin-streptomycin Mixture (Lonza), and in a 37 °C incubator with a humidified atmosphere of 5% CO2 in air. Cells were infected with HSV-1 at an MOI of 1, then they were incubated for 1 h. The virus suspension was removed and cells were washed with PBS. This step was followed by the addition of fresh medium to the cells and they were incubated for 1, 2, 4, 6, 8, or 12 h. A mixture from the different time points were prepared for the further analysis.

##### BoHV-1

Madin–Darby Bovine Kidney (MDBK) cells were used for the infection using the Cooper isolate (GenBank Accession # JX898220.1) of Bovine Herpesvirus 1.1. Cells were incubated at 37 ◦C in a humidified incubator with 5% CO2 and were cultured in DMEM supplemented with 5% (v/v) fetal bovine serum, 100 U/mL penicillin, and 100 µg/mL streptomycin. Cells were infected using MOI=1 virus suspension. Infection was allowed to progress until complete cytopathic effect was observed. Infected cells were incubated for 1, 2, 4, 6, 8, and 12 h, then the supernatant was collected, and the cellular fraction was subjected to three successive cycles of freezing and thawing in order to release additional intracellular virions.

### RNA isolation

#### Extraction of total RNA

Total RNA was isolated from the PRV, EHV-1, VZV and KSHV infected cells by using the NucleoSpin® RNA kit (Macherey-Nagel). The spin-column protocol was applied. In brief, cells were lysed by the addition of a chaotropic ion containing buffer solution (from the kit). Nucleic acids were then bound to a silica membrane. Samples were treated with DNase I to remove genomic DNA. Total RNAs were eluted with RNase-free water. To eliminate the potential remaining DNA from the samples, we used the TURBO DNA-free™ Kit. Samples were stored at -80 °C.

#### Purification of polyadenylated RNA

The Qiagen Oligotex mRNA Mini Kit was used to enrich the mRNAs (and other RNAs with polyA-tail) from the PRV, EHV-1 and VZV samples, which were then used as templates for ONT and Illumina library preparations. The Spin Columns protocol of the manual was applied. In short, the final volume of the RNA samples was set to 250 µL by the addition of RNase-free water. Then, 15 µL Oligotex suspension and 250 µL OBB buffer (both from the Oligotex kit) were added to the mixtures, which were first incubated at 70°C for 3 min and then at 25 °C for 10 min. Samples were centrifuged at 14,000×g for 2 min, then supernatants were removed. Four-hundred µL Oligotex OW2 wash buffer was added to the samples, then they were spun down in Oligotex spin columns at 14,000×g for 1 min. This step was repeated once, and finally, the poly(A)+ RNA fraction was eluted from the membrane by adding 60 µl hot Oligotex elution buffer. In order to maximize the yield, a second elution step was also carried out. The Lexogen Poly(A) RNA Selection Kit V1.5 was used for the selection of polyadenylated RNAs from the KSHV total RNA samples. Briefly, 10 µL of total RNA (5 µg) was denatured at 60°C for 1 min then it was hold at 25°C. The RNA samples were mixed with 10 µL washed bead (including in the kit) and the mixtures were incubated at 25°C for 20 min with 1,250 rpm agitation. Afterwards, the tubes were placed in a magnetic rack for 5 min. Supernatant was discarded and the beads were resuspended in Bead Wash Buffer (supplied by the Lexogen kit) and were incubated 25°C for 5 min with 1,250 rpm agitation. Tubes were transferred onto the magnetic rack and supernatant was discarded after 5 min incubation. This washing step was carried out twice. After the second washing step, the beads were resuspended in Nuclease-free water (Lexogen kit) and then incubated at 70°C for 1 min. Then, the tubes were transferred onto the magnet for 5 min. The supernatant containing the poly(A)+ RNA fraction was transferred into a fresh tube.

#### Removal of rRNA

Ribo-Zero Magnetic Kit H/M/R (Epicentre/Illumina) was used to remove ribosomal RNAs and to enrich mRNAs. Unlike poly(A)+ purification, the rRNA depletion process also retains RNAs without polyA tails, except for rRNAs. For starting material, 5 µg of a mixture of EHV-1 total RNA was used. The sample was mixed with the Ribo-Zero Reaction Buffer and Ribo-Zero rRNA Removal Solution. The mixture was incubated at 68°C for 10 min, then at room temperature for 5 min. Next, 225 µl washed Magnetic Bead was added to the sample and they were incubated at room temperature for 5 min, then at 50°C for 5 min. Finally, the mixture was placed on a magnetic stand, then the supernatant containing the rRNA-depleted RNA was collected. The final purification was carried out by using the Agencourt RNAClean XP Beads (Beckman Coulter) as recommended by the Ribo-Zero manual.

#### Enrichment of the 5′ ends of RNAs

Terminator™ 5′-Phosphate-Dependent Exonuclease (Lucigen) was used to enrich the 5′ ends of the transcripts. The process was carried out with a mixture of poly(A)+ RNAs from the EHV-1 samples, which was mixed with Terminator 10X Reaction Buffer A, RiboGuard RNase Inhibitor and Terminator Exonuclease (1 Unit). The mixture was incubated at 30°C for 60 min, then the reaction was stopped by the addition of 1 µL of 100 mM EDTA (pH 8.0). RNAClean XP beads (Beckman Coulter) was used for final purification.

### Long-read sequencing

#### Direct cDNA sequencing

Libraries were generated without PCR amplification from the poly(A)+ fractions of RNAs from PRV, EHV-1 and KSHV as well as from the Terminator-handled EHV-1 samples. For this approach, the ONT’s Direct cDNA Sequencing Kit (SQK-DCS109) was used according to the ONT’s manual. Briefly, the RNAs were mixed with ONT VN primer and 10 mM dNTPs, and incubated at 65°C for 5 min. Next, RNaseOUT (Thermo Fisher Scientific), 5x RT Buffer (Thermo Fisher Scientific), ONT Strand-Switching Primer were added to the mixtures and they were incubated at 42°C for 2 min. Maxima H Minus Reverse Transcriptase enzyme (Thermo Fisher Scientific) was added to the samples to generate the first cDNA strands. The reaction was carried out at 42°C for 90 min, and the reactions were terminated by heating the samples at 85°C for 5 min. The RNAs from the RNA:cDNA hybrids were removed by using RNase Cocktail Enzyme Mix (Thermo Fisher Scientific). This reaction was performed at 37°C for 10 min. The LongAmp Taq Master Mix [New England Biolabs (NEB)] and ONT PR2 Primer were used to synthetize the second strand of cDNAs. The following PCR condition was applied: 1 min at 94 °C, 1 min 50°C, then 15 min at 65°C. As a next step, the end-repair and dA-tailing were carried out with the NEBNext End repair /dA-tailing Module (NEB) reagents. These reactions were performed at 20°C for 5 min and then the samples were heated at 65°C for 5 min. This was followed by the adapter ligation using the NEB Blunt /TA Ligase Master Mix (NEB) at room temperature for 10 minutes. ONT Native Barcoding (12) Kit was used to label the libraries, then they were loaded to ONT R9.4.1 SpotON Flow Cells (200 fmol mixture of libraries was loaded to one flow cell. AMPure XP Beads were used after each enzymatic step. Samples were eluted in UltraPure™ nuclease-free water (Invitrogen).

##### Amplified Nanopore cDNA sequencing

For more accurate mapping of the 5′-end of VZV transcripts, random-primer based RT reactions were carried out as a first step of library preparation. For this, poly(A)-selected and Terminator-treated RNA samples and SuperScript IV enzyme (Life Technologies) were used. From the first-strand cDNAs, libraries were prepared according to the modified 1D strand switching cDNA by ligation protocol Ligation Sequencing kit (SQK-LSK108; Oxford Nanopore Technologies). KAPA HiFi DNA Polymerase (Kapa Biosystems) and Ligation Sequencing Kit Primer Mix (part of the 1D Kit) were used to amplify the cDNAs. Next, samples were end-repaired using NEBNext End repair / dA-tailing Module (New England Biolabs), then adapter ligation was carried out using the sequencing adapters supplied with the kit and NEB Blunt/TA Ligase Master Mix (New England Biolabs).

##### Direct RNA sequencing

Direct RNA sequencing (dRNA-seq) approach was also applied for library preparation from the EHV-1and KSHV samples with the aim to avoid potential biases associated with reverse transcription (RT) and PCR reactions to detect and validate novel splice variants as well as 3′ UTR isoforms. For the dRNA-seq experiments, we used an RNA mixture containing RNA from each time points of infection from the Poly(A)+ samples and from the Poly(A)+ and Terminator-treated RNAs. The oligo dT-containing T10 adapter for RT priming and the RNA CS for monitoring the sequencing quality (both from the ONT kit) were added to the RNA mix along with NEBNext Quick Ligation Reaction Buffer, and T4 DNA ligase (both from NEB). The reaction was incubated for 10 min at room temperature. Then, 5 × first-strand buffer, DTT (both from Invitrogen), dNTPs (NEB) and UltraPure™ DNase/RNase-Free water (Invitrogen) were added to the samples. Finally, SuperScript III enzyme (Thermo Fisher Scientific) was mixed with the sample and RT reaction was performed at 50°C for 50 min, and subsequently stopped by heat inactivation of the enzyme at 70°C for 10 min. RNA adapter (from the ONT kit) was ligated to the RNA:cDNA hybrid sample using the NEBNext Quick Ligation Reaction Buffer and T4 DNA ligase at room temperature for 10 min. The RNAClean XP Beads were used after each additional enzymatic step. Two flow cells were used for dRNA-seq, 100 fmol from the sample was loaded onto each of them.

### Short-read sequencing

For the SRS transcriptomic analysis of EHV-1 virus, libraries were generated from a mixture of poly(A)+ enriched and rRNA-depleted samples using the NEXTflex® Rapid Directional qRNA-Seq Kit (PerkinElmer). Fragmentation of the RNAs was carried out enzymatically at 95°C for 10 min via the addition of NEXTflex® RNA Fragmentation Buffer. Next, the first strand cDNA was synthetized. First, the NEXTflex® First Strand Synthesis Primer was mixed with the RNA sample, which were incubated at 65 °C for 5 min, then the mixture was subsequently placed on ice. RT reaction was performed by using the NEXTflex® Directional First Strand Synthesis Buffer and Rapid Reverse Transcriptase according the following protocol: incubation at 25°C for 10 min, then at 50°C for 50 min, and termination at 72°C for 15 min. Generation of the second cDNA strand was carried out at 16°C for 60 min with the addition of NEXTflex® Directional Second Strand Synthesis Mix (with dUTPs). Adenylation of cDNAs were performed at 37°C for 30 min using the NEXTflex® Adenylation Mix. The reaction was terminated by incubating the samples at 70°C for 5 min. As a next step, Molecular Index Adapters (part of the Kit) were ligated to the sample using the NEXTflex® Ligation Mix. Ligation was carried out at 30°C for 10 min. Samples were amplified by PCR. As a first step of this, the NEXTflex® Uracil DNA Glycosylase was added to the samples, and then they were incubated at 37°C for 30 min, which was followed by heating at 98°C for 2 min, and they were subsequently placed on ice. Then, the following components were added to the samples: PCR Master Mix, qRNA-Seq Universal forward primer, and qRNA-Seq Barcoded Primer (sequence: AACGCCAT; all from the PerkinElmer kit). The following settings were applied for the PCR: 2 min at 98 °C, followed by 15 cycles of 98 °C for 30 sec, 65 °C for 30 sec and 72 °C for 60 sec, followed by a final extension at 72 °C for 4 min. AMPure XP Bead was used after each enzymatic reaction. The NEXTflex® Resuspension buffer was used for the final elution of the sequencing library from which 10 pM was loaded to the Illumina MiSeq reagent cassette. Paired-end transcriptome sequencing was performed on an Illumina MiSeq sequencer by using the MiSeq Reagent Kit v2 (300 cycles).

### Measurement of nucleic acid quality and quantity

#### RNA

The Qubit 4. fluorometer and the Qubit Assay Kits (Invitrogen) were used for the measurement of RNA concentration. The RNA BR Assay was applied for the quantitation of total RNA samples whereas the RNA HS Assay was utilized for the poly(A)+ and ribodepleted RNA fractions. The quality of the total RNA samples was checked by using the Agilent TapeStation 4150 device, RNA ScreenTape and reagents were used. RIN scores above 9.6 were used for cDNA production. The RNA quality was assessed with the Agilent 2100 Bioanalyzer (for PacBio sequencing) or Agilent 4150 TapeStation System (for MinION sequencing) and RIN scores above 9.6 were used for cDNA production.

#### cDNA

The Qubit dsDNA HS Assay Kit (Invitrogen) was used to quantify the cDNA samples. For the analysis of Illumina library quality, the Agilent High Sensitivity D1000 ScreenTape was used.

### Real-time RT-PCR

Quantitative RT2-PCR was used for transcript validation and kinetic analysis, and was carried out as previously described^103^ using the primers listed in **Supplementary Table 3.** Briefly, first strand cDNAs from the total RNAs of PRV, EHV-1, VZV, HSV-1, BoHV-1 and KSHV were synthesized by using the SuperScript III reverse transcriptase and gene-specific primers (ordered from Integrated DNA Technologies). The reactions were carried out in 5μl of final volume containing 0.1 μg of total RNA, 2 pmol of the primer, dNTP mix (Invitrogen, 10 μM final concentration), 5x First-Strand Buffer, and the RT enzyme (Invitrogen). The reaction was performed at 55 °C for 60 min and then terminated by heating the samples to 70 °C for 15 min. Ten-fold dilutions from the RT products were used as template for real-time PCR amplification. PCR reactions were carried out using ABsolute QPCR SYBR Green Mix (Thermo Fisher Scientific) in a Rotor-Gene Q (Qiagen) cycler. The running conditions are shown in **Supplementary Table 4**. The following controls were used: loading control (28S rRNA as a reference gene), no template (checking the potential primer-dimer formation) and no RT control (to detect the potential DNA contamination). The relative expression levels (R) were calculated using the mathematical model described by Soong and colleagues^104^ with the following modifications: the R values were calculated using the average E^Ct^ value of the 6h samples for each gene as a control, which was normalized with the average of the corresponding 28S values (E^Ct-^reference). We used the Comparative Quantitation module of the Rotor-Gene Q software package that automatically calculates the qPCR efficiency sample-by-sample and set the cycling thresholds values automatically.

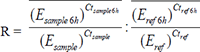

### Cap Analysis of Gene Expression (CAGE)

We applied CAGE-Seq to uncover the TSS distribution patterns at the examined genomic regions in EHV-1 and KSHV using three biological replicates. The CAGE™ Preparation Kit (DNAFORM, Japan) was applied. CAGE libraries were prepared from 5 µg of total RNA, in three replicates, following the manufacturer’s recommendations. Briefly, the RNA and the RT primer (random primer mixture from CAGE™ Prep Kit) was denatured at 65 °C for 5 min. First cDNA strands were synthesized by using SuperScript III Reverse Transcriptase (Invitrogen). In order to increase the activity and specificity of RT enzyme, trehalose/sorbitol mixture (CAGE™ Prep Kit) was also added. The reactions were incubated for 30 sec at 25 °C, then the RT reaction was carried out at 50 °C for 60 min. Diol groups in the Cap at the 5′-end (and ribose at 3′-end) were oxidized with NaIO_4_ and Biotin (long arm) hydrazine was bound to it. First, the oxidation was carried out with the addition of NaOAc (1M, pH 4.5, CAGE™ Prep Kit) and NaIO_4_ (250mM, CAGE™ Prep Kit) to the samples and incubation on ice for 45 min in the dark. After this step, 40% glycerol and Tris-HCl (1M, pH 8.5, CAGE™ Prep Kit) were added to the samples. NaOAc (1M, pH 6.0) and Biotin Hydrazine (10 mM, CAGE™ Prep Kit) were mixed with the samples and the oxidized diol residues were biotinylated at 23 °C for 2 hours. After this step, single strand RNA was digested by applying RNase I (CAGE™ Prep Kit) treatment (37 °C 30 min). Biotinylated, Capped RNA samples were mixed and bound (Cap-trapping) to the pretreated Streptavidin beads (pretreatment details below) at 37 °C 30 min. Then they were incubated on a magnetic rack. The beads were washed with Wash Buffer 1 (twice), then with Wash Buffer 2, and finally with Wash Buffer 3 (both from the CAGE™ Prep Kit). Next, cDNAs were released from the beads: Releasing Buffer was added to the samples and they were incubated at 95 °C for 5 min. After a short incubation on a magnetic rack, the supernatant (containing the Capped cDNAs) was transferred to new tubes. RNase I buffer (CAGE™ Prep Kit) was added to the tRNA-Streptavidin bead and they were placed on a magnetic rack. The supernatant was transferred to the tubes containing the cDNAs and they were stored on ice. The samples were treated with an RNase mixture (RNase H and RNase I, both from the CAGE™ Prep Kit) and incubated at 37 °C for 15 min. The potential remaining RNA was digested with RNase I. The reaction was performed at 37 °C for 30 min.

Streptavidin beads were coated with tRNA (CAGE™ Prep Kit), which was followed by mixing and incubation on ice for 30 min, and then incubated on a magnetic stand for 3 min. Supernatant was removed, and the beads were washed with Wash Buffer 1 (CAGE™ Prep Kit) twice. Finally, the beads were eluted in Wash Buffer 1 and tRNA was also added. The volume of the samples was reduced by using the miVac DUO Centrifugal Concentrator (Genevac), then single strand 5′ linkers (with barcodes) were ligated to the samples at 16 °C for 16 hours using the DNA ligation mixture CAGE™ Prep Kit). After a purification step, the miVac DUO was used again in order to concentrate the samples. This was followed by the ligation of the 3′ linker using the DNA ligation mixture. It was carried out at 16 °C for 16 hours. (The 5′ and 3′ linkers were pre-heated at 55 °C and the cDNA samples at 95 °C before the ligation steps). The samples were mixed with Shrimp Alkaline Phosphatase (SAP, CAGE™ Prep Kit) to remove the phosphate group of the linkers. The reaction was conducted at 37 °C for 30 min and terminated at 65 °C for 15 min. Then USER enzyme was added to the sample, which digests the dUTP from 3′ linker up strand. The USER treatment was carried out at 37 °C for 30 min and it was stopped at 95 °C for 5 min. After this step the barcoded samples were mixed and concentrated with miVac DUO. Finally, the second cDNA strands were synthesized with the 2nd primer, DNA polymerase, buffer and 10 mM dNTP (all from the CAGE™ Prep Kit). The denaturation step was set to 95 °C for 5 min, the annealing to 55 °C for 5 min, and the elongation to 72 °C for 30 min. The sample mixture was treated with Exonuclease I enzyme (37 °C for 30 min). The vacuum concentrator was used to completely dry the sample and finally it was dispensed in 10 µl of nuclease-free H2O. The amount of single-stranded cDNAs was measured using Qubit 2.0 and the Qubit ssDNA HS Assay Kit. The RNAClean XP Beads were used after RT, oxidation and biotinylation. AmpureXP Beads were used to purify the samples after Cap-trapping and Releasing, RNase I treatment, 5′ and 3′ linker ligation, SAP and USER treatments, 2nd strand cDNA synthesis and Exonuclease I treatment. Libraries with different barcodes were pooled and applied on the same flow cells. The libraries were sequenced on a MiSeq instrument with v3 (150 cycles) and v2 (300 cycles) chemistries (Illumina). Qubit 4.0 and 1X dsDNA High Sensitivity (HS) Assay was used to measure the concentration of the sample. The quality of the library was tested by TapeStation.

### Bioinformatic analyses

### Illumina CAGE sequencing data analysis

Read quality was checked with **fastqc** (https://www.bioinformatics.babraham.ac.uk/projects/fastqc). The reads were trimmed with **TrimGalore** (https://github.com/FelixKrueger/TrimGalore) using the following settings: *-length 151 -q*. The **STAR** aligner^105^, version 2.7.3.a was used to map the reads to the KSHV strain TREx reference genome (*GQ994935.1*) using *--genomeSAindexNbases 8,* otherwise default parameters. The **CAGEfightR** R package^106^ was used to identify TSSs and TSS clusters using a minimum pooled value cutoff of 0.1 (*pooledcutoff=0.1*).

### ONT sequencing data analysis

**Guppy** software (v3.4.5) was used to basecall the ONT-MinION sequencing reads. Reads passing the quality filter of 8 (default), were mapped to the reference genome using **minimap2**^107^ with the following settings: -*ax splice -Y -C5 -cs.* **ReadStatistics** script from **Seqtools** (https://github.com/moldovannorbert/seqtools) was used to compute mapping statistics. The **LoRTIA** toolkit (alpha version, accessed on 20 August 2019, https://github.com/zsolt-balazs/LoRTIA) was used to identify TESs, TSSs and introns and to subsequently reconstruct transcripts using these features.

The LoRTIA workflow with default settings was as follows: 1.) for dRNA and dcDNA sequencing: *−5 TGCCATTAGGCCGGG --five_score 16 -- check_in_soft 15 −3 AAAAAAAAAAAAAAA --three_score 16 s Poisson–f true*; and 2.) for o(dT)-primed cDNA reads: *−5 GCTGATATTGCTGGG -- five_score 16 -- check_in_soft 15 −3 AAAAAAAAAAAAAAA --three_score 16 s Poisson–f true*.

In case of EHV-1, the searching for adapter sequences was performed by the following command: samprocessor.py --five_adapter GCTGATATTGCTGGG --five_score 14 --check_in_soft 15 --three_adapter AAAAAAAAAAAAAAAAAAA --three_score 14 input output. The next step in the workflow was the annotation of TSS and TES. For the TES positions, the wobble has been increased to 20, the TSS wobble value remains the default. Then following parameters were used on the ‘sam’ files: Stats.py -r genome -f r5 -b 10 and Stats.py -r genome -f l5 -b 10 for the TSS detection; while Stats.py -r genome -f r3 -b 20 and Stats.py -r genome -f l3 -b 20 for TES detection; and Stats.py -r genome -f in for intron detection.

Several sequencing techniques were used in the analysis of BoHV-1, the LoRTIA workflow was set up differently depending on sequencing techniques. The following parameters were used for dRNA sequencing: *LoRTIA -5 AGAGTACATGGG --five_score 16 --check_in_soft 15 -3 AAAAAAAAAAAAAAAAAAAAA -- three_score 12 -s poisson -f True*. For oligo d(T) cDNA sequencing: *LoRTIA -5 TGCCATTAGGCCGGGGG -- five_score 14 --check_in_soft 15 -3 AAAAAAAAAAAAAAAAAAAAA --three_score 14 -s poisson -f True.* During random primer cDNA sequencing: *LoRTIA -5 TGCCATTAGGCCGGGGG --five_score 14 --check_in_soft 15 - 3 GAAGATAGAGAGCGACA --three_score 14 -s poisson -f True.* Also for dcDNA sequencing: *LoRTIA -5 GCTGATATTATTGCTGGG --five_score 16 --check_in_soft 15 -3 AAAAAAAAAAAAAAAAAAAAAAA -- three_score 14 -s poisson -f True*

A TSS was accepted if the adapters were correct while TESs were accepted if polyA tails were present and there were no false priming events detected by LoRTIA. Further, in the case of KSHV, EBV, EHV-1, TSS were accepted only if at least one dcDNA read, and one dRNA or CAGE read validated it. In the case of CMV, BoHV-1, PRV and HSV-1, TSSs were accepted only if they were present in at least three different samples. In case of introns, we accepted only those that were present in the dRNA sequencing, as this method is considered to be the ‘*Gold Standard*’ in identifying alternative splicing events. Several transcripts were manually included if they were a long TSS variant of already accepted TSSs. **MotifFinder** from **Seqtools** was used to find promoter elements around the accepted TSSs.

### Downstream data analysis and visualization

Downstream data analysis was carried out within the R environment, using **GenomicRanges**^108^, **tidygenomics**^109^ and packages from the **tidyverse**^110^. **Gviz**^111^ was used to create **Figures 2-4** and **Supplementary Figures 2-7** (https://github.com/ivanek/Gviz). **Figure 5** was created using a custom R program. Briefly, the ‘.bam’ files were imported into R using **Rsamtools** (https://bioconductor.org/packages/Rsamtools) then the 5′ ends were summed per genomic position and finally a density plot was generated using **ggplot2’s** geom_density geom function, using the following parameters: adjust = 0.025 but otherwise default parameters (including the default Gaussian kernel). The density along with the genome annotation was plotted using a custom plotting function, utilizing gggenes (https://github.com/wilkox/gggenes). These scripts can be used to import other alignments into R and to generate similar plots from their 3′ or 5′ distributions on the reference genomes as well. These scripts area available on GitHub at https://github.com/Balays/R.codes

#### Accession codes

The LoRTIA software suite and the SeqTools are available on GitHub.

LoRTIA: https://github.com/zsolt-balazs/LoRTIA.

SeqTools: https://github.com/moldovannorbert/seqtools.

R.codes: https://github.com/Balays/R.codes

## Acknowledgements

This research was supported by National Research, Development and Innovation Office (NRDIO), Researcher-initiated research projects (Grant numbers: K 128247 and K 142674) to ZB and by the NRDIO Research projects initiated by young researchers (Grant number: FK 128252) to DT. The work was also supported by NIH grant R01AI132554 to ZT. The APC fee was covered by the University of Szeged, Open Access Fund: 5954.

## Ethics declarations

### Author contributions

**GT**: carried out bioinformatics, data analysis, interpretation and integration of data

**DT**: carried out library preparation, long-read – and CAGE sequencing, participated in interpretation of data and drafted the manuscript

**AIAA**: carried out RNA purification and long-read sequencing, participated in analysis

**ZC**: carried out viral infection, propagation of cells, RNA purification, long-read sequencing and qPCR

**GÁN**: participated in bioinformatics and interpretation of data

**BK**: participated in bioinformatics and interpretation of data

**GG**: participated in sequencing and bioinformatics

**LMS**: participated in propagation of cells, carried out RNA purification

**IG**: participated in data analysis

**ÁF**: participated in bioinformatics and interpretation of data

**ÁD**: participated in RNA purification, long-read sequencing

**IP**: participated in RNA purification and data analysis

**MM**: carried out propagation of cell cultures, long-read participated in sequencing

**VÉD**: participated in propagation of cells and interpretation of data

**VC**: participated in propagation of cells

**ZZ**: propagated cells and viruses, participated in RNA purification

**ZT**: participated in the design of the experiments and drafted the manuscript

**ZB**: conceived and designed the experiments, supervised the study, wrote the manuscript All authors read and approved the final paper.

## Abbreviations

αHV: alphaherpesvirus
asRNA: antisense RNA
βHV: betaherpesvirus
BoHV-1: Bovine alphaherpesvirus 1
CDS: coding sequence
DBP: DNA-binding protein
DNP: DNA polymerase
dRNA-Seq: direct RNA sequencing
dcDNA-Seq: direct cDNA sequencing
E: early
EBV: Epstein-Barr virus
EHV-1: Equid alphaherpesvirus
γHV: gammaherpesvirus
HCMV: Human cytomegalovirus
HHV-6: Human herpesvirus 6
HSV-1: Herpes simplex virus 1
IE: immediate-early
ICP: infected cell polypeptide
IR: inverted repeat
IRL: internal repeat of UL region
IRS: internal repeat of US region
L: late
LAT: latency-associated transcript
LLT: long latency transcript
lncRNA: long noncoding
RNA LRS: long-read sequencing
L/ST: L/S junction-spanning transcript
miRNA: micro RNA
ncRNA: non-coding RNA
ONT: Oxford Nanopore Technologies
ORC: origin recognition complex
ORF: open reading frame
Ori: replication origin
PacBio: Pacific Biosciences
PRV: Pseudorabies virus
raRNA: replication origin-associated
RNA RNP: RNA polymerase
SRS: short-read sequencing
SVV: Simian varicella virus
TES: transcript end site
TF: transcription factor
TI: transcript isoform
TO: transcriptional overlap
TR: transcription regulator
TRL: terminal repeat of UL region
TRS: terminal repeat of US region
TSS: transcript start site
UL: unique long
ES: unique short
UTR: untranslated region
VZV: Varicella-zoster virus

## Supplementary Figures

**Suplementary Figure 1. Location of replication origins on the herpesvirus genomes**

This figure shows the location of inverted repeats and replication origins in the genomes of herpesviruses examined in this study. *Abbreviations* Ul: unique long, US: unique short, IRS: internal repeat of US region, TRS: terminal repeat of US region, IRL: internal repeat of UL region, TRL: terminal repeat of UL region

**Supplementary Figure 2. Ori-proximal transcripts of BoHV-1**

This picture shows the transcripts encoded by the OriS-proximal region of Bovine herpesvirus 1 (IRS: A; TRS: B). All of the putative transcripts were identified by LoRTIA software using dcDNA datasets, unless indicated otherwise. Protein coding genes are labeled with black arrows, non-coding genes with green arrows, mRNAs with blue arrows, and ncRNAs with red arrows. The relative transcript abundance was labeled by shades. The coverage of dRNA-Seq data in BoHV-1 is relative low therefore, undetection certain RNAs with this technique does not necessarily mean low reliability. Transcripts that have a proximal TATA box are marked by a ‘T’ letter, and those that were also detected by dRNA-Seq are marked with a ‘d’ letter at the upstream positions. The vertical red arrows indicate the positions of TATA boxes on the genome. Introns are indicated by horizontal lines.

**Supplementary Figure 3. Ori-proximal transcripts of PRV**

This picture shows the transcripts encoded by the OriL-(A) and OriS-proximal regions of Pseudorabies virus. All the putative transcripts were identified by LoRTIA software using dcDNA datasets, unless indicated otherwise. Protein coding genes are labeled with black arrows, non-coding genes with green arrows, mRNAs with blue arrows, and ncRNAs with red arrows. The relative transcript abundance was labeled by shades. Those transcripts that have a proximal TATA box are marked by a ‘T’ letter, and the vertical red arrows indicate the positions of TATA boxes on the genome. Those transcripts that were also detected by dRNA-Seq are marked with a ‘d’ letter at the upstream positions. Introns are indicated by horizontal lines.

**Supplementary Figure 4. Ori-proximal transcripts of SVV**

This picture shows the transcripts encoded by the OriS-proximal region of Simian varicella virus. All of the putative transcripts were identified by LoRTIA software using dcDNA datasets, unless indicated otherwise. Protein coding genes are labeled with black arrows, non-coding genes with green arrows, mRNAs with blue arrows, and ncRNAs with red arrows. The relative transcript abundance was labeled by shades. The vertical red arrows indicate the positions of TATA boxes on the genome.

**Supplementary Figure 5. Ori-proximal transcripts of HSV-1**

This picture shows the transcripts encoded by the Ori-proximal regions (OriL: A; OriS of IRS: B; OriS of TRS: C) of Herpes simplex virus 1. All the putative transcripts were identified by LoRTIA software using dcDNA datasets, unless indicated otherwise. Protein coding genes are labeled with black arrows, non-coding genes with green arrows, mRNAs with blue arrows, and ncRNAs with red arrows. The relative transcript abundance was labeled by shades. Those transcripts that have a proximal TATA box are marked by a ‘T’ letter, and the vertical red arrows indicate the positions of TATA boxes on the genome. Those transcripts that were also detected by dRNA-Seq are marked with a ‘d’ letter at the upstream positions. The striped ends of certain arrows (illustration of transcripts) indicate that these terminals have been not or not accurately annotated.

**Supplementary Figure 6. Ori-proximal transcripts of HCMV**

This picture shows the transcripts specified by the Ori-proximal region of Human cytomegalovirus. All the putative transcripts were identified by LoRTIA software using dcDNA datasets, unless indicated otherwise. Protein coding genes are labeled with black arrows, non-coding genes with green arrows, mRNAs with blue arrows, and ncRNAs with red arrows. The relative transcript abundance was labeled by shades. The vertical red arrows indicate the positions of TATA boxes on the genome.

**Supplementary Figure 7. Ori-proximal transcripts of EHV**

This picture shows the transcripts specified by the Ori-proximal regions of Epstein-Barr virus (OriP: A; Orilyt-L: B; OriLyt-R: C). All of the putative transcripts were identified by LoRTIA software using dcDNA datasets, unless indicated otherwise. Protein coding genes are labeled with black arrows, non-coding genes with green arrows, mRNAs with blue arrows, and ncRNAs with red arrows. The relative transcript abundance was labeled by shades. The vertical red arrows indicate the positions of TATA boxes on the genome. In the case of very long transcripts, the names of the genes overlapped by these transcripts are enlisted. Introns are indicated by horizontal lines.

**Supplementary Figure 8. The operation of the Putative Super Regulatory Center**

A. Passage of RNA polymerases across the OriS by reading the long 5′ UTR of *us1* and *icp4* genes inhibits the assembly of ORC and replisome. Stalling of RNP on OriS leads to the same consequences.

B. Reading the long 5′ UTRs of *us1* and *icp4* genes by RNP inhibits each other expressions which exert an effect on the global transcription. Inhibition of *icp4* expression through interference between RNPs reading the two genes leads a decreased expression in most of the herpesvirus genes, including *us1* gene. Inhibition of *us1* gene expression ICP22 inhibits *icp4* expression and leads a more complex effect on genome-wide gene expression (Takács et al., 2013).

C. Transcriptional overlapping of replication origin may lead to the formation of RNA:DNA hybrids and thereby interfering (inhibiting or perhaps facilitating) the assembly of the replisome.

D. Transcriptional overlapping of replication origin and the transcription regulatory genes may lead to the formation of RNA:DNA hybrids and thereby interfering (inhibiting or perhaps facilitating) the assembly of the replisome and also the transcription.

**Supplementary Figure 9. Interference between the replication and transcription apparatuses**

A. Passage of RNA polymerases across the OriS by reading the long 5′ UTR of *ul29* and *ul30* genes inhibits the assembly of ORC and replisome. Stalling of RNP on OriS leads to the same consequences.

B. Head-on collision between the two replication genes leads to the inhibition of each other transcription and the initiation replication because of the passage of RNPs across the OriL. The replication is also affected by the inhibition of the generation of replication proteins.

C. Formation of RNA:DNA hybrids at the OriL interferes with the initiation of DNA replication.

D. Formation of RNA:DNA hybrids at the OriL and the replication genes interferes with both the initiation of DNA replication and the transcription of replication genes. The replication is also affected by the inhibition of the generation of replication proteins.

## Supplementary Tables

**Supplementary Table 1. Correspondence of orthologous transcripts**

**Supplementary Table 2. Transcripts with putative regulatory functions**

**Supplementary Table 3. List of the primers used for RT^2^-PCR**

**Supplementary Table 4. Running conditions of RT^2^-PCR**

The annealing temperature was increased to 65°C for the amplification of PRV ELIE, HSV-1 ICP4-OriS and KSHV LIR1-ORF71 transcripts.

