## Supplementary figures and images for "Novel Herpesvirus Transcripts with Putative Regulatory Roles in DNA Replication and Global Transcription"

### Supplementary Figure

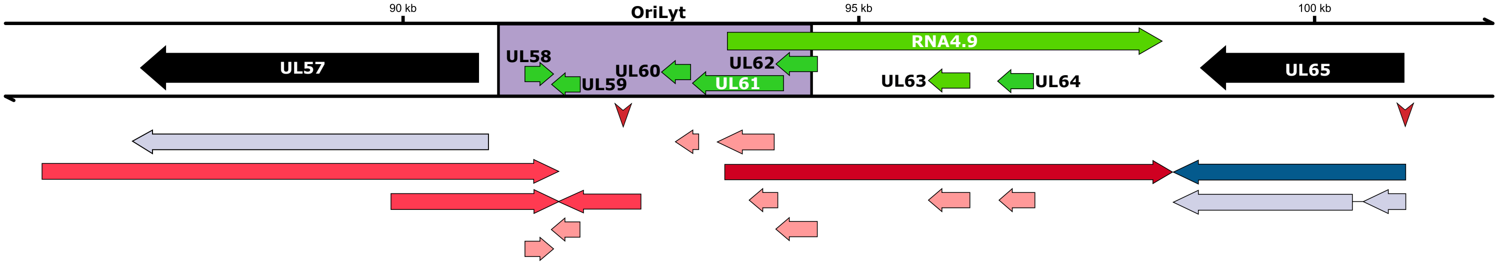

### Supplementary Figure 1

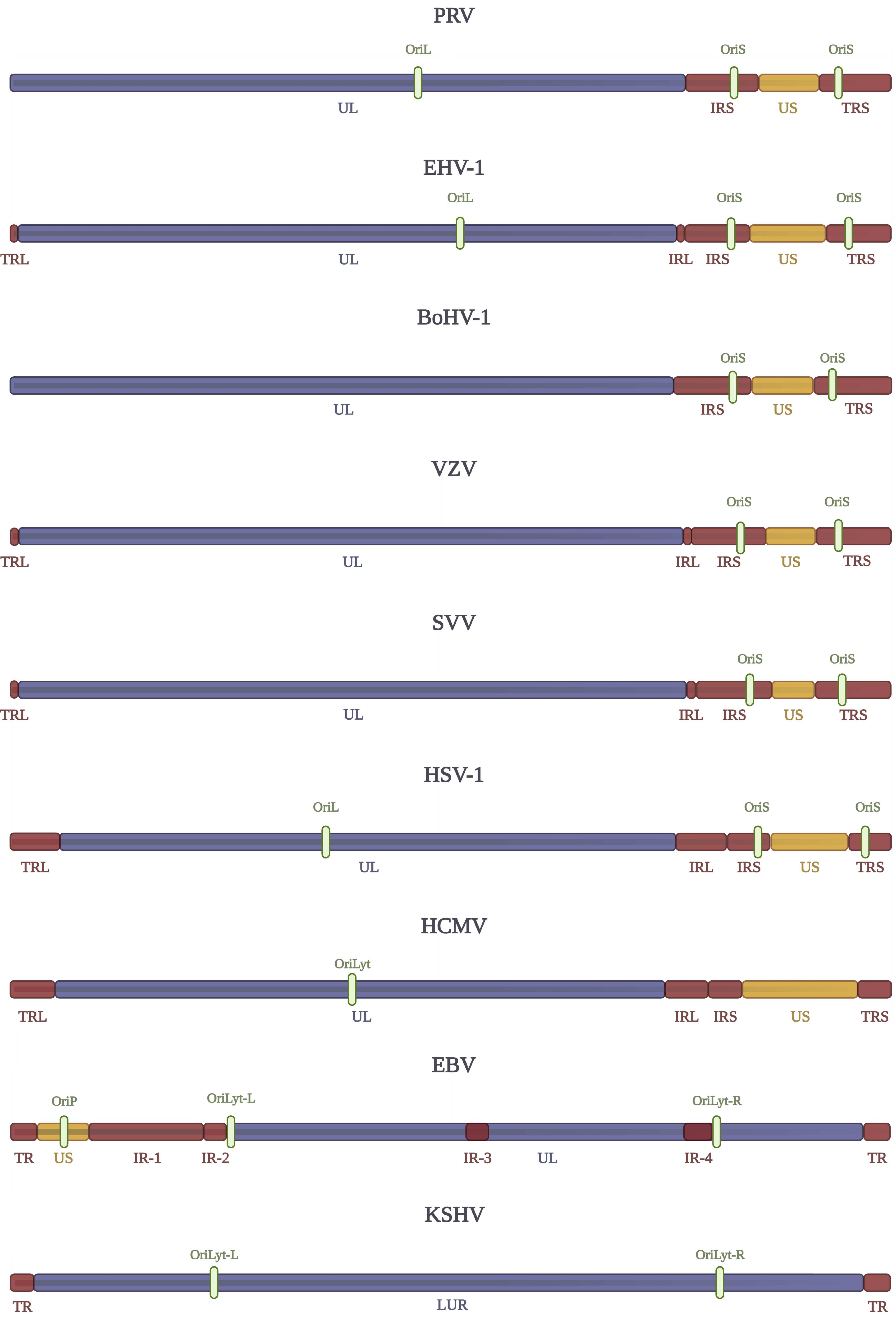

### Supplementary Figure 2

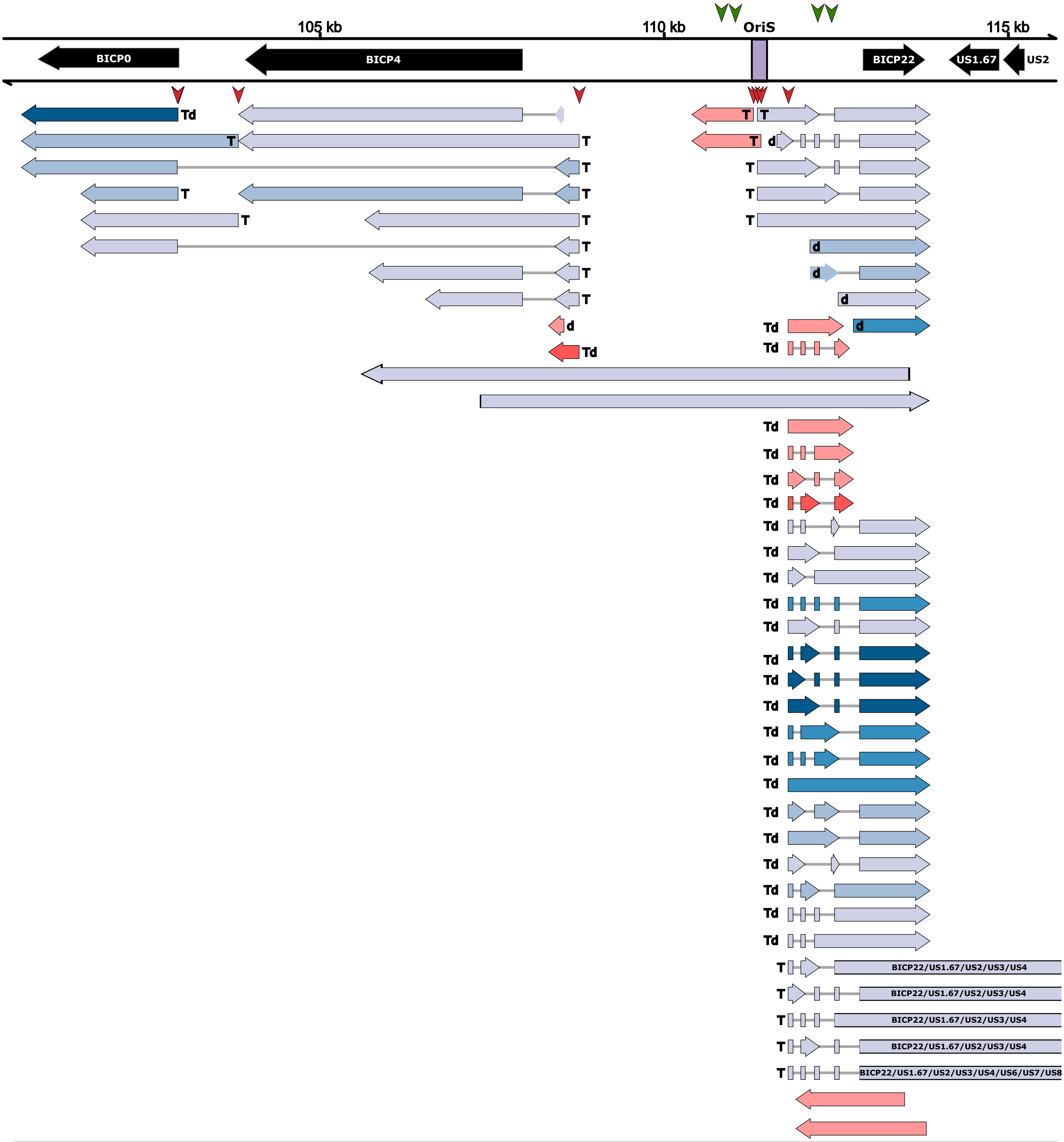

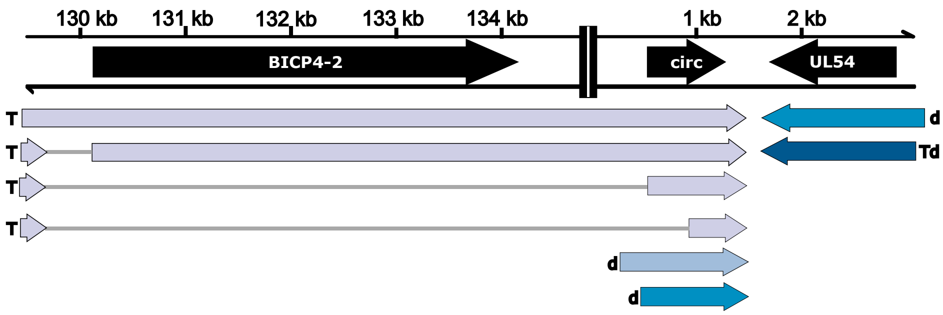

### Supplementary Figure 3

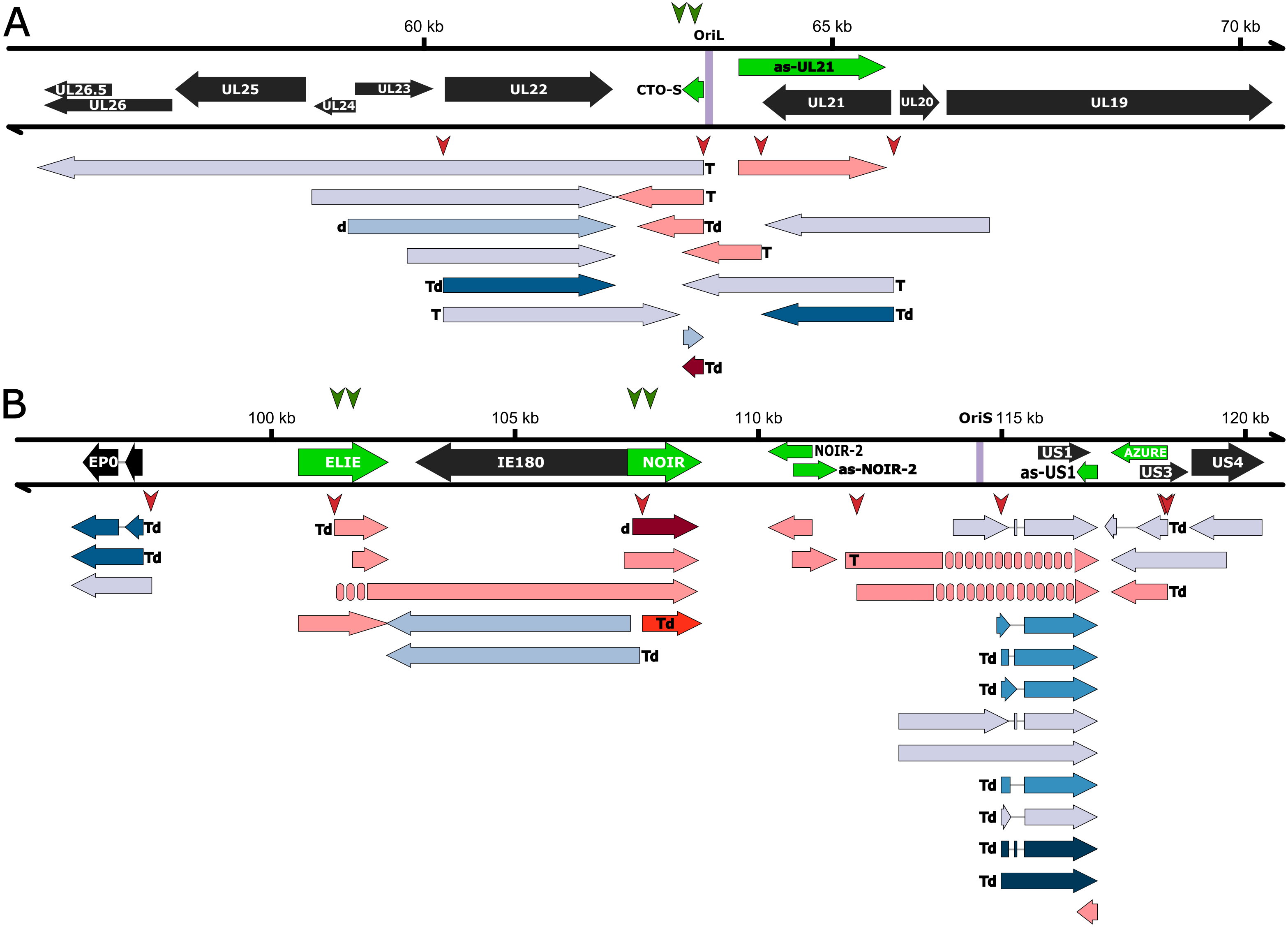

### Supplementary Figure 4

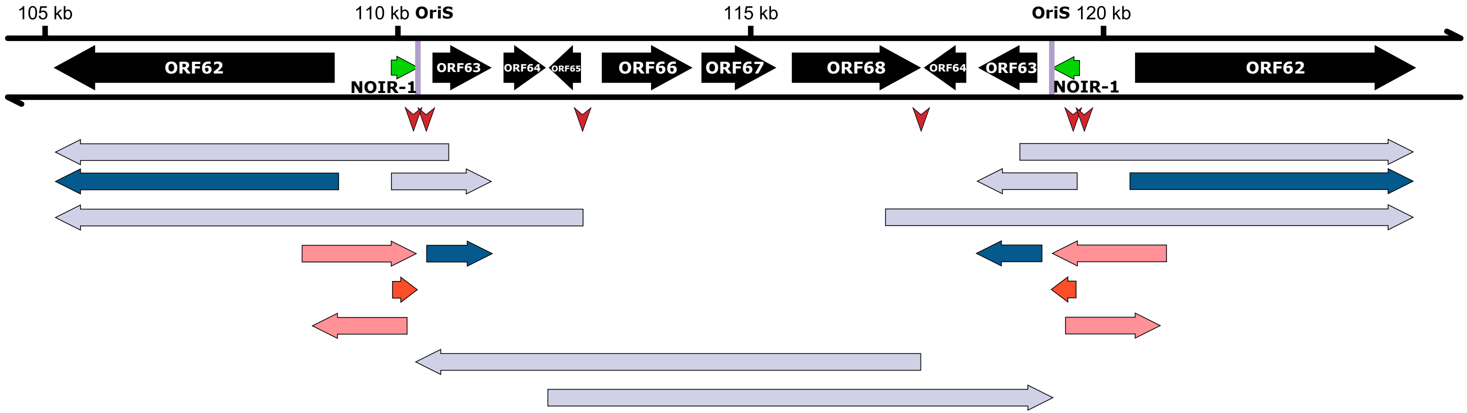

### Supplementary Figure 5

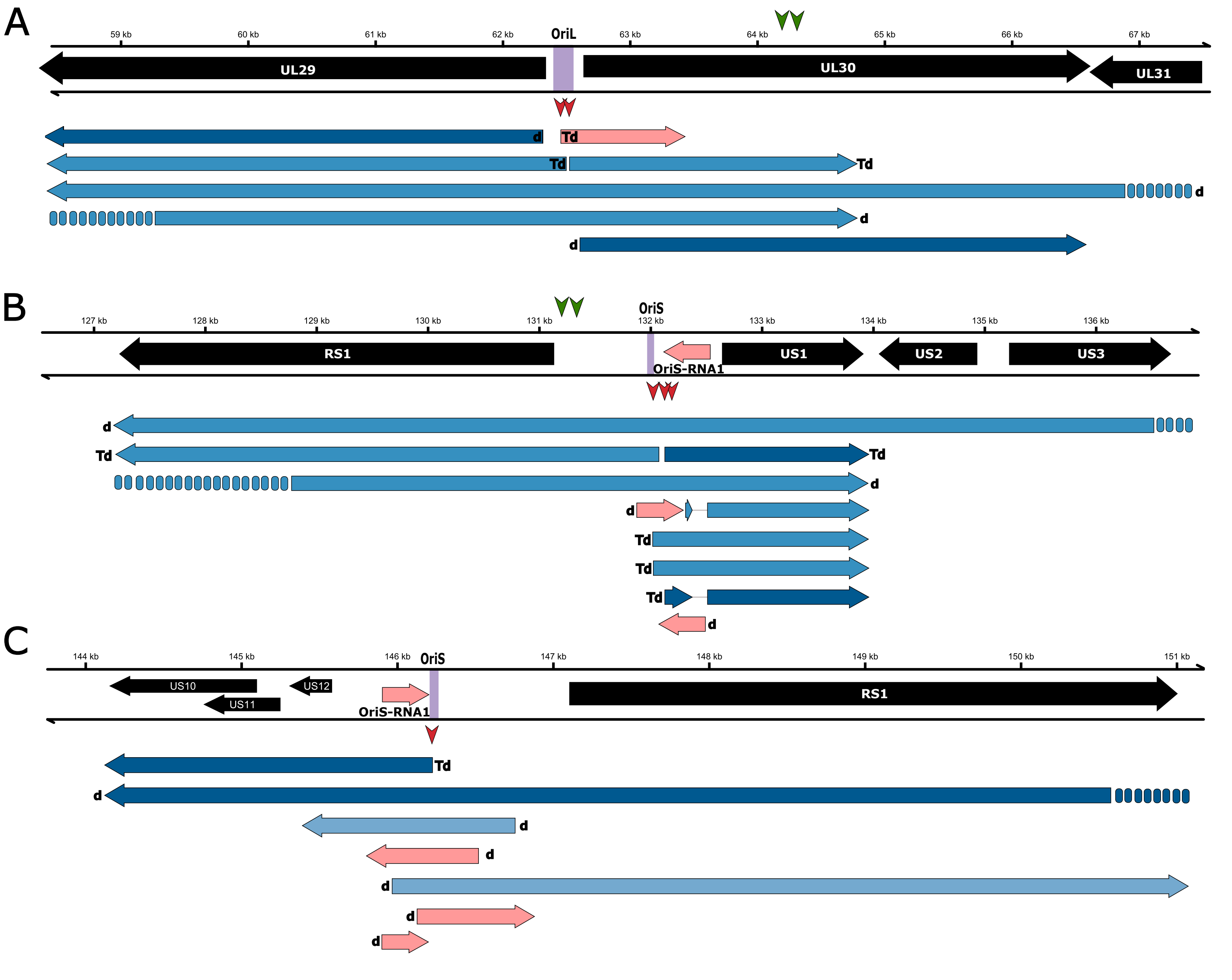

### Supplementary Figure 7

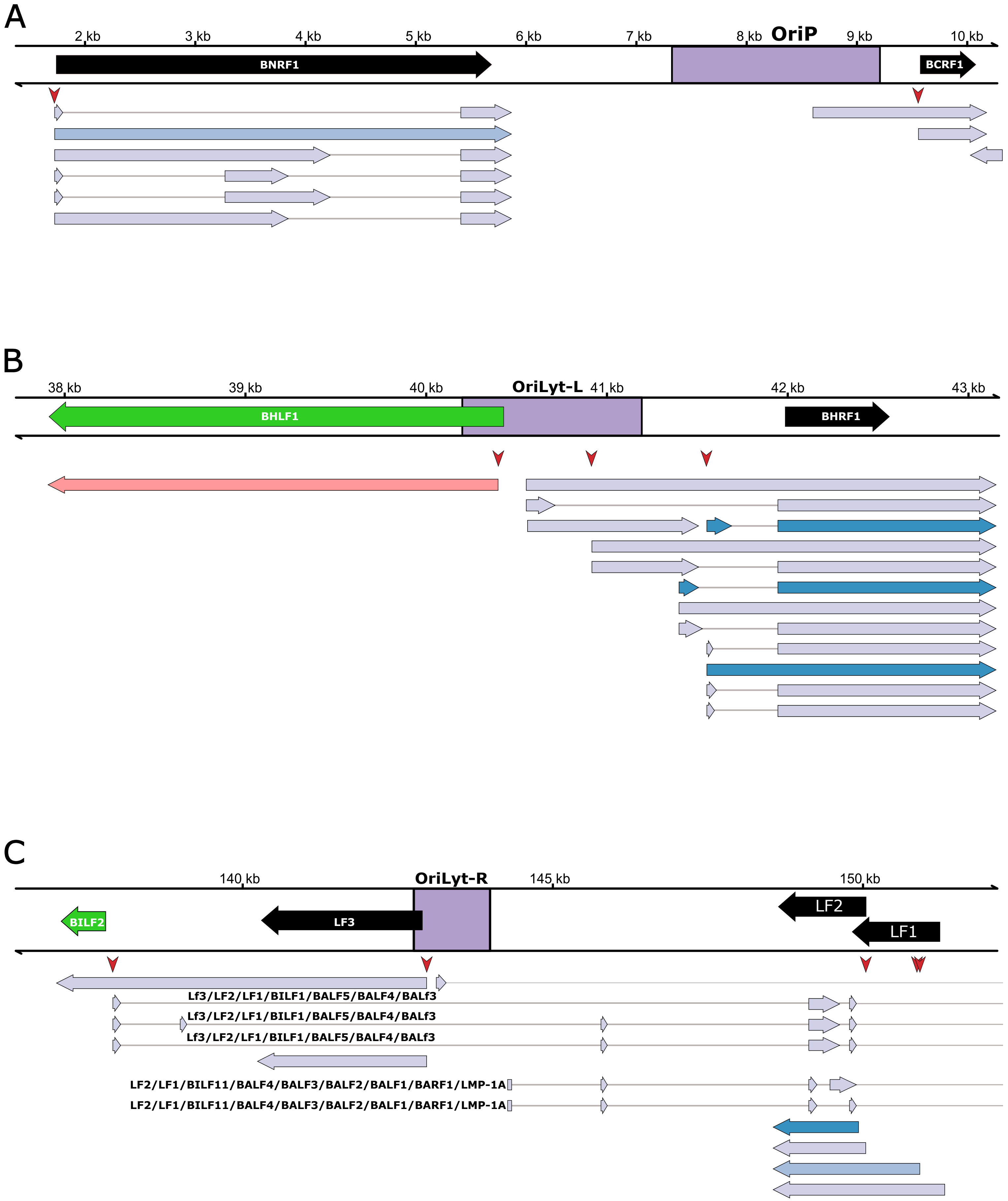

### Supplementary Figure 8

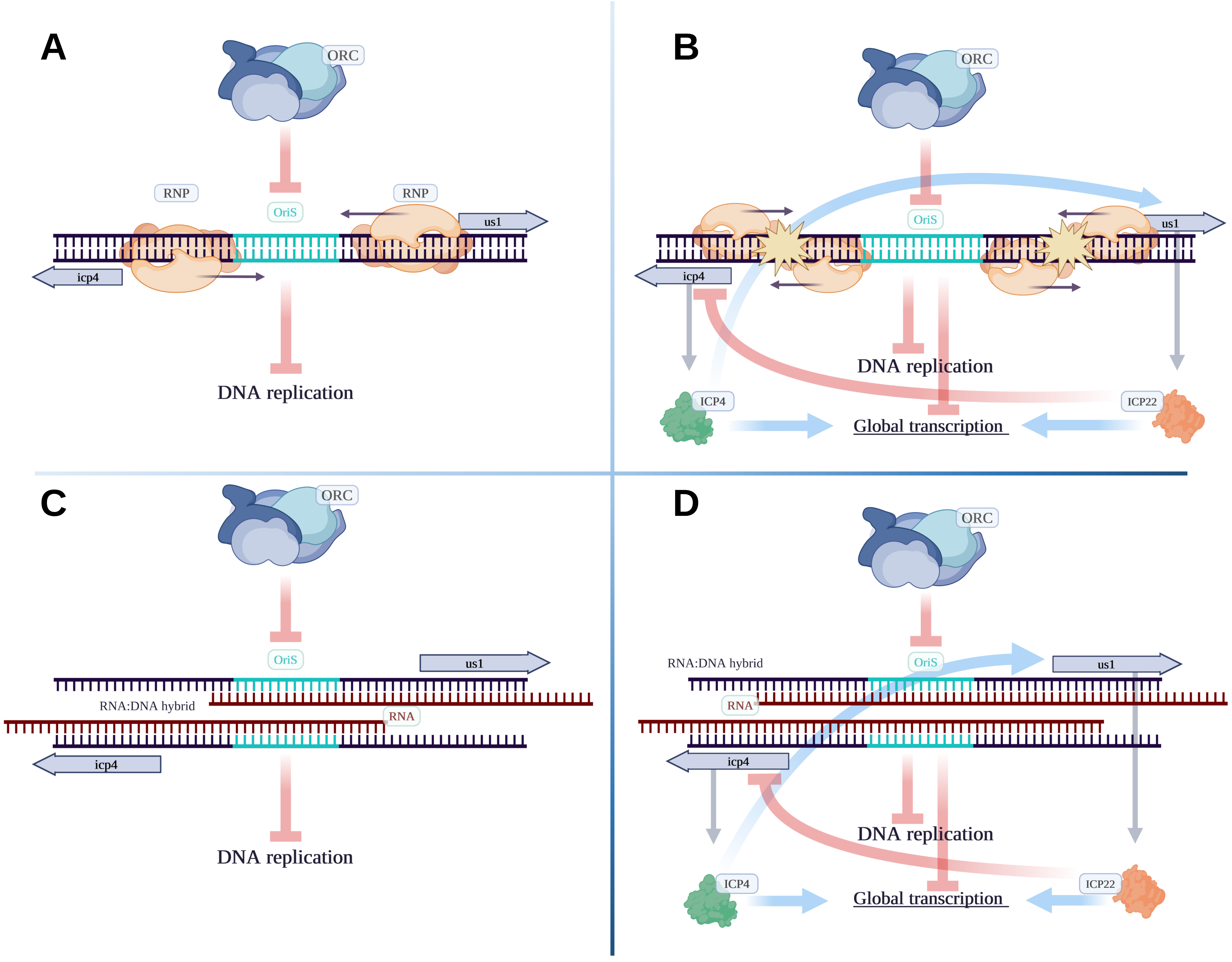

### Supplementary Figure 9

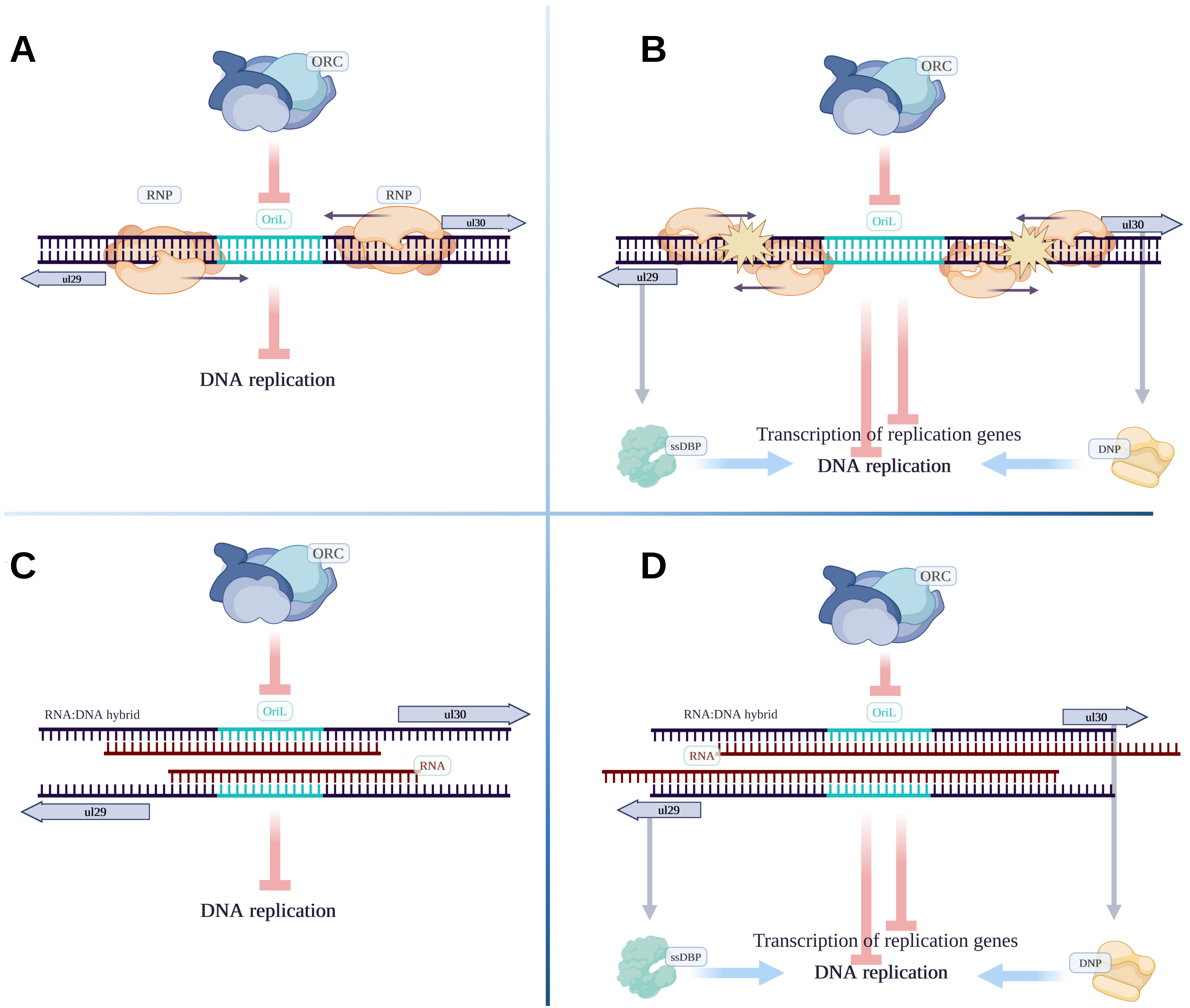
