## Supplementary Table 4 for "Novel Herpesvirus Transcripts with Putative Regulatory Roles in DNA Replication and Global Transcription"

**Supplementary Table 4. Running conditions of RT^2^-PCR**

|  |  | Temperature | Time | Cycle number |
| --- | --- | --- | --- | --- |
| pre-denaturation |  | 95°C | 15 min | 1 |
| denaturation |  | 94°C | 25 sec | 30 |
| annealing |  | 60°C * | 25 sec |  |
| extension |  | 72°C | 6 sec |  |
